# Mesophyll conductance response to short-term changes in CO_2_ is related to leaf anatomy and biochemistry in diverse C_4_ grasses

**DOI:** 10.1101/2021.10.03.462792

**Authors:** Varsha S. Pathare, Robert J. DiMario, Nouria Koteyeva, Asaph B. Cousins

## Abstract

- Mesophyll CO_2_ conductance (g_m_) in C_3_ species responds to short-term (minutes) changes in environment potentially due to changes in some leaf anatomical and biochemical properties and due to measurement artifacts. Compared to C_3_ species, there is less information about g_m_ responses to short-term changes in environment conditions like *p*CO_2_ across diverse C_4_ species and the potential determinants of these responses.
- Using 16 diverse C_4_ grasses we investigated the response of g_m_ to short-term changes in CO_2_ and how this response related to the leaf anatomical and biochemical traits.
- For all the measured C_4_-grasses g_m_ increased as CO_2_ decreased; however, the percent change in g_m_ varied (+13% to +250%) and significantly related to percent changes in leaf transpiration efficiency (TE_i_). The percent increase in g_m_ was highest in grasses with thinner mesophyll cell walls and greater leaf nitrogen, activities of phosphoenolpyruvate carboxylase (PEPC), Rubisco and carbonic anhydrase, and a higher affinity of PEPC for bicarbonate.
- Our study demonstrates that CO_2_ response of g_m_ varies greatly across diverse C_4_ grasses and identifies the key leaf anatomical and biochemical traits related to this variation. These findings have implications for improving C_4_ photosynthetic models, and in attempts to improve TE_i_ through manipulation of g_m_.

## Introduction

Mesophyll CO_2_ conductance (g_m_) describes the movement of CO_2_ from substomatal cavities across the intercellular air space, cell walls and membranes to the site of first carboxylation. This carboxylation occurs in the mesophyll chloroplast in species with C_3_ photosynthetic pathway and mesophyll cytosol in species with C_4_ photosynthetic pathway (Evans & von Caemmerer, 1996). g_m_ varies across plant groups due to variation in leaf anatomy and biochemistry, changes dynamically in response to environmental stimuli and has a significant impact on the plant and ecosystem-level photosynthetic CO_2_ uptake and water-use efficiency (Evans *et al*., 2009; Flexas *et al*., 2014; Knauer *et al*., 2019b; Pathare *et al*., 2020a). Despite its importance for photosynthetic CO_2_ uptake and water-use at both plant and ecosystem levels and its variation across diverse plants groups, g_m_ is only beginning to be explicitly implemented into global models which upscale leaf-scale photosynthetic processes to canopy and global scales (Rogers *et al*., 2017; Knauer *et al*., 2019a; Knauer *et al*., 2019b; von Caemmerer, 2021).The implementation of g_m_ is complicated because there is a lack of detailed information about responses of g_m_ to short and long-term changes in environmental conditions (e.g., light, precipitation, temperature, and CO_2_ concentration) across diverse plant groups (Rogers *et al*., 2017; Knauer *et al*., 2019a; Knauer *et al*., 2019b). Most investigations on the responses of g_m_ to changes in environmental conditions focus on C_3_ species (von Caemmerer & Evans, 2015; Xiong *et al*., 2015; Carriqui *et al*., 2018; Shrestha *et al*., 2018), but there is less information about the response of g_m_ to changing environmental conditions in diverse C_4_ species (Ubierna *et al*., 2017; Kolbe & Cousins, 2018; Ubierna *et al*., 2018; Sonawane *et al*., 2021). A better understanding of how C_4_-g_m_ responds to changing environmental conditions is essential for improving the models of C_4_ photosynthetic at both the leaf and global scales (Rogers *et al*., 2017; Knauer *et al*., 2019a; Knauer *et al*., 2019b; von Caemmerer, 2021) as well as for potentially increasing water-use efficiency of C_4_ crops through manipulation of g_m_ (Cousins *et al*., 2020; Pathare *et al*., 2020a).

The influence of g_m_ on C_3_ photosynthesis has been well studied for the past several years and there has been a significant understanding about C_3_-g_m_ and its responses to changes in long-term growth conditions and short-term measurements conditions. In general, g_m_ varies greatly among diverse C_3_ species and limits C_3_ photosynthesis as much as g_sw_ (Flexas *et al*., 2014; Muir *et al*., 2014; Barbour & Kaiser, 2016; Veromann-Jürgenson *et al*., 2017). Variation in C_3_-g_m_ is influenced by leaf ontogenic and anatomical traits like leaf development and senescence (Grassi & Magnani, 2005; Barbour *et al*., 2016), surface area of chloroplasts appressed to intercellular air space (Sc)(Tosens *et al*., 2012; Peguero-Pina *et al*., 2016), mesophyll cell wall thickness (M_CW_) (Veromann-Jürgenson *et al*., 2017; Ellsworth *et al*., 2018; Evans, 2021) and leaf thickness (Flexas *et al*., 2008; Muir *et al*., 2014). In terms of responses to long-term growth conditions, C_3_-g_m_ is influenced by water stress, elevated CO_2_, salinity, nutrient supplement, and growth latitude (Flexas *et al*., 2008; Momayyezi & Guy, 2017; Mizokami *et al*., 2018; Shrestha *et al*., 2018). Whereas rapid responses of g_m_ (within minutes) have been observed in response to short-term changes in leaf temperature, quantity and quality of light, relative humidity, and CO_2_ concentrations (Hassiotou *et al*., 2009; von Caemmerer & Evans, 2015; Xiong *et al*., 2015), though not always (Loreto *et al*., 1992; Tazoe *et al*., 2009). However, there is no consensus on the exact cause of rapid responses of g_m_ to changes in environmental conditions like CO_2_.

Some studies suggest that rapid changes in g_m_ in C_3_ plants can be explained by changes in chloroplast position and movement that could lead to short-term change in S_c_ (Oguchi *et al*., 2005; Tholen *et al*., 2008; Terashima *et al*., 2011). Whereas mesophyll cell wall thickness (T_CW_) and composition are considered invariable in short-term (minutes) and cannot explain the rapid responses of g_m_ (Evans *et al*., 2009; Terashima *et al*., 2011; Carriqui *et al*., 2018). Alternatively, the rapid responses of C_3_-g_m_ have also been attributed to changes in activities of key photosynthetic enzymes like carbonic anhydrase (CA)-which catalyzes the conversion of CO_2_ to HCO_3_^-^ (Evans *et al*., 2009; Momayyezi & Guy, 2017) and the facilitation effect of CO_2-_ permeable aquaporins (Uehlein *et al*., 2008; Flexas *et al*., 2012; Kaldenhoff, 2012; Groszmann *et al*., 2017), but see (Kromdijk *et al*., 2019). Still, others suggest that rapid responses of C_3_-g_m_ could be the result of systematic methodological errors or the use of oversimplified models. These include mathematical dependency of g_m_ on other variables (like A_net_ and C_i_) used to calculate it in the fluorescence method, neglecting the contribution of respiratory and photorespiratory CO_2_ release to the total CO_2_ pool in the leaf and inaccurate measurement of day respiration or estimates of the Rubisco fractionation factor in the Δ^13^C method (Tholen & Zhu, 2011; Gu & Sun, 2014; Carriqui *et al*., 2018; Ubierna *et al*., 2019). The exact mechanism underlying the CO_2_ responses of C_3_-g_m_ thus remains controversial. However, considerable evidence based on diverse species and methods of estimating g_m_ suggest that C_3_-g_m_ increases at low *p*CO_2_.

There has been a recent increase in research on the short- and long-term variability of g_m_ in diverse C_4_ species and the leaf traits that could explain this variability (Barbour *et al*., 2016; Cano *et al*., 2019; Pathare *et al*., 2020a; Pathare *et al*., 2020b). Our recent study demonstrated that, like C_3_ species, g_m_ varies significantly among diverse C_4_ grasses and has significant effects on photosynthetic rates and leaf water-use efficiencies (Pathare *et al*., 2020a). This variation in C_4_-g_m_ was correlated with leaf-level traits like leaf thickness, stomatal ratio, adaxial stomatal densities and S_mes_. We also demonstrated that C_4_ grasses adapted to low precipitation habitats exhibit traits related to greater g_m_ but lower leaf hydraulic conductance compared to grasses from habitats with relatively high precipitation (Pathare *et al*., 2020b). These studies have advanced our understanding of the variability of g_m_ in C_4_ grasses as well as how g_m_ is influenced by leaf-level traits and is affected by long-term growth conditions like precipitation. However, there is still a limited understanding of how g_m_ in diverse C_4_ grasses responds to short-term changes in environmental conditions like CO_2_. The few previous studies have largely focused on a few species like sorghum, maize and *Setaria* (Osborn *et al*., 2017; Kolbe & Cousins, 2018; Ubierna *et al*., 2018; Sonawane & Cousins, 2020). To the best of our knowledge, no studies to date have explored if g_m_ responses to short-term changes in *p*CO_2_ vary among diverse C_4_ grasses and which potential anatomical and biochemical traits could explain this variation in CO_2_ response of C_4_-g_m_.

The overall objectives of the current study were (1) to investigate the response of g_m_ to short-term changes in *p*CO_2_ in diverse C_4_ species, (2) identify the anatomical and biochemical traits that may explain the variable CO_2_ response of C_4_-g_m_, and (3) evaluate the impact of varying CO_2_ responses of g_m_ on changes in photosynthesis and leaf water-use efficiency. To address above objectives, we estimated g_m_ under changing *p*CO_2_ in 16 diverse C_4_ grasses (see Pathare *et al*., 2020a for details). Several methods have been used to estimate C_4_-g_m_. Pfeffer & Peisker (1998) calculated g_m_ from the initial slope of a photosynthetic CO_2_ response curve and assumed no CO_2_ dependence of g_m_. However, C_4_-g_m_ is sensitive to *p*CO_2_ (Kolbe & Cousins, 2018; Ubierna *et al*., 2018) and hence the initial slope method may be problematic. Using anatomical traits like S_mes_ for estimating C_4_-g_m_ could also be subjected to errors due to assumptions made for values of T_CW_, porosity, and membrane permeability (Pengelly *et al*., 2010). The Δ^13^C *in vitro* V_pmax_ method (Ubierna *et al*., 2017) estimates C_4_-g_m_ by retrofitting models of C_4_ photosynthesis model (von Caemmerer, 2000) and the Δ^13^C (Farquhar & Cernusak, 2012) with gas exchange, kinetic constants and *in vitro* PEPC activities. The Δ^13^C *in vitro* V_pmax_ method may also result in inaccurate estimates of C_4_-g_m_ due to errors associated with estimation of *in vitro* PEPC and CA activities and enzyme kinetic parameters. The Δ^18^O method (Gillon & Yakir, 2000; Barbour *et al*., 2016) utilizes simultaneous measurements of the oxygen isotope composition (δ^18^O) of transpired H_2_O and oxygen isotope discrimination (Δ^18^O) in CO_2_ to calculate the CO_2_ concentration at the site of isotope exchange by CA. The Δ^18^O method assumes full isotopic equilibrium between CO_2_ and H_2_O at the site of CA (θ = 1), which may not always be true and hence g_m_ could be underestimated (Barbour *et al*., 2016). In the current study we used the method described by (Ogee *et al*., 2018) that builds upon the Δ^18^O method by accounting for the possibility for an incomplete CO_2_-H_2_O equilibrium inside leaves, the physical separation between mesophyll and bundle sheath cells in C_4_ leaves and the contribution of respiratory fluxes.

We also investigated the relationship of leaf anatomical traits, previously known to influence C_4_-g_m_ (Pathare *et al*., 2020a), with variable CO_2_ response of g_m_ in the 16 C_4_ grasses. Furthermore, we investigated the impacts of photosynthetic capacity on the CO_2_ response of g_m_ in these C_4_ grasses, through measurements of maximum photosynthetic rates (A_max_), leaf nitrogen content (N_area_), activities of key enzymes of C_4_ photosynthetic pathway like CA, phosphoenolpyruvate carboxylase (PEPC) and ribulose-1,5-bisphosphate carboxylase/oxygenase (Rubisco) and PEPC’s affinity for its substrate (HCO^-^_3_). N_area_ is integral to the proteins of photosynthetic machinery (like PEPC, Rubisco and CA) which, along with the leaf structure, are responsible for drawdown of CO_2_ inside the leaf (Parkhurst, 1994; Wright *et al*., 2004; Evans *et al*., 2009; DiMario *et al*., 2018).

## Material and methods

### Plant growth

Sixteen C_4_ grasses (Table 1) were selected for this study. In the graphs, each species is identified by four letter word which combines the first letter of genera and first three letters of species name. CO_2_ response of g_m_ was not measured for two C_4_ grasses, *Sporobolus airoides* Torr. and *Echinochloa esculenta* (A.Braun) H. Scholz that have been included in our previous studies (Pathare *et al*., 2020a; Pathare *et al*., 2020b). Details about growth conditions have been described in our previous study (Pathare *et al*., 2020a). In short, plants were raised from seeds and grown in 3-L free drainage pots in a controlled environment growth chamber (model GC-16; Enconair Ecological Chambers Inc., Winnipeg, MB, Canada). The photoperiod was 14 h including a 2 h ramp at the beginning and end of the light period. Light and dark temperatures were maintained at 26 and 22 °C, respectively. Light was provided by 400-W metal halide and high-pressure sodium lamps with maximum photosynthetic photon flux density (PPFD) of ca. 1000 μmol photons m^-2^ s^-1^ at plant height. One individual per species was grown per pot in a Sunshine mix LC-1 soil (Sun Gro Horticulture, Agawam, MA, USA) with 5-6 replicate pots per species. The plants were irrigated daily to pot saturation and fertilized twice a week with Peters 20-20-20 (2.5 g L^-1^). Pots were randomized daily within the growth chamber.

**Table 1:**
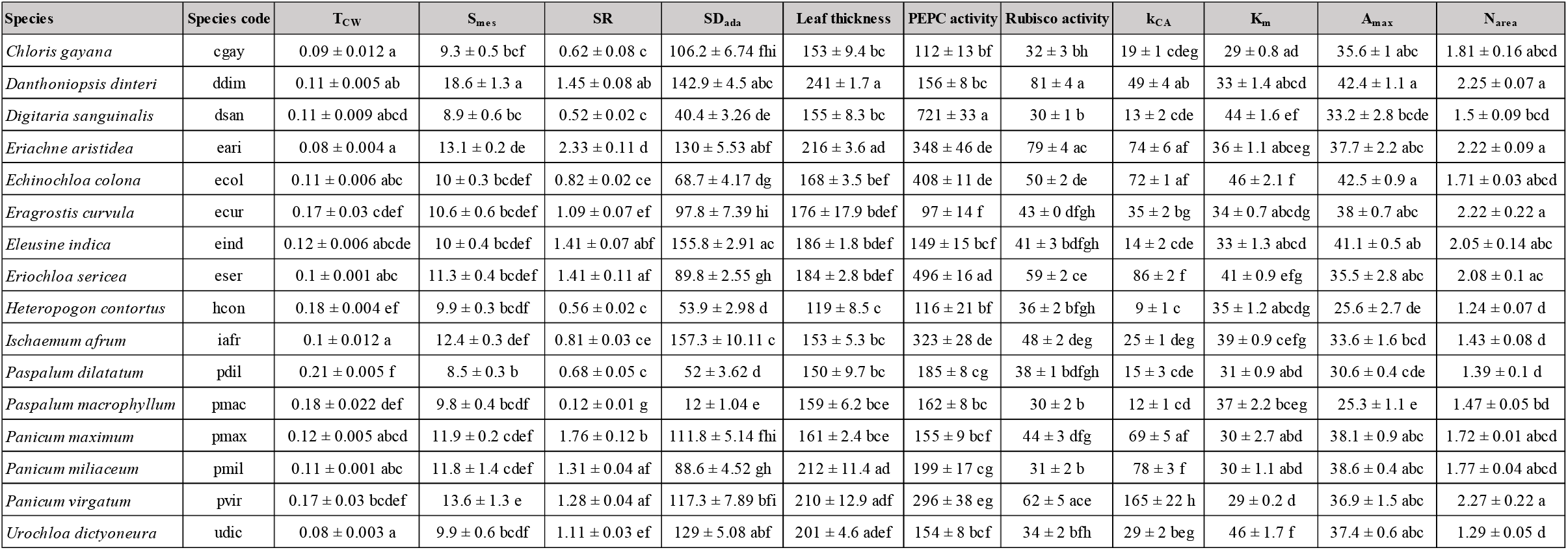
Mean ± SE (*n* = 3 to 6) values along with the corresponding letters of post-hoc Tukey’s test for important leaf-level anatomical and biochemical traits measured for 16 C_4_ grasses. Species code was created using first letter of genera and first three letters of species. Results of one-way ANOVA are given in Table S1. Values for T_CW_, S_mes_, SR, SD_ada_, Leaf thickness, k_CA_ and N_area_ have been published in Pathare *et al*., 2020a, whereas values for K_m_ have been published in DiMario *et al*., 2021.

### Gas exchange measurements and estimation of g_m_

The measurements of net photosynthetic rates (A_net_, μmol CO_2_ m^-2^ s^-1^), stomatal conductance to water vapor (g_sw_, μmol water m^-2^ s^-1^), intercellular CO_2_ concentrations (C_i_, Pa), and mesophyll conductance to CO_2_ (g_m_, μmol m^-2^ s^-1^ Pa^-1^) were performed (Pathare *et al*., 2020a) at four different CO_2_ levels inside chamber (CO_2_S) -34, 27, 20 and 14 Pa. Intrinsic water-use efficiency (TE_i_, μmol CO_2_ mol^-1^ water) was calculated as A_net_/g_sw_ for each CO_2_ level. Maximum photosynthetic rates (A_max_, μmol CO_2_ m^-2^ s^-1^) were measured at saturating light of ∼1200 μmol photons m^-2^ s^-1^ and *p*CO_2_ of ∼1500 μmol m^-2^ s^-1^. For estimating g_m_, isotopologs of CO_2_ and H_2_O were measured using the LI-6400XT infrared gas analyzer (LiCor, Lincoln, NE, USA) coupled to a tunable diode laser absorption spectroscope (TDLAS, model TGA 200A, Campbell Scientific, Logan, UT, USA) and a cavity-ring down absorption spectroscope (Picarro, Sunnyvale, CA, USA) as described previously (Ubierna *et al*., 2017; Kolbe & Cousins, 2018; Pathare *et al*., 2020a). The entire LI6400XT, the 2 cm × 6 cm leaf chamber (6400-11, Li-Cor), and LI-6400-18-RGB light source were placed in a growth cabinet (model EF7, Conviron; Controlled Environments Inc., MN, USA) with fluorescent lamps (F48T12/CW/ VHO; Sylvania, Wilmington, MA, USA) set at a PPFD of ∼250 μmol photons m^-2^ s^-1^ and air temperature was maintained at 25 °C. g_m_ was estimated using the most recent method described by Ogee *et al*., (2018) as discussed in Pathare *et al*., (2020a). This method utilizes a newly developed model of C_4_ photosynthetic discrimination that provides an estimate of the isotopic equilibration between CO_2_ and H_2_O inside the leaf and g_m_. The CO_2_ responses of g_m_, g_sw_, A_net_ and TE_i_ were measured by maintaining a mole fraction of CO_2_ in the leaf chamber. For each species, four-point CO_2_ response curve was performed. During measurements CO_2_ sample (CO_2_S) was set to about 34, 27, 20 and 14 Pa. During measurement at each CO_2_ level, the leaves were allowed to adjust for atleast 30 minutes or until stable values of A_net_ and g_sw_ were achieved. Data were subsequently collected and averaged over the next 20-30 min and the Li6400XT was set to log data only when the TDLAS analyzed the sample line. Three to six biological replicates were measured per species.

Percent change in key physiological traits like A_net_, g_sw_, g_m_ and TE_i_ in response to decrease in CO_2_S (from 34 to 14 Pa) was calculated as : (mean trait value at 14 Pa-mean trait value at 34 Pa)*100/ (mean trait value at 34 Pa)

### Measurement of anatomical traits and habitat mean annual precipitation

Light and electron microscopy techniques were used to measure important structural and anatomical traits like adaxial stomatal density (SD_ada_, number mm^-2^), abaxial stomatal density (SD_aba_, number mm^-2^), stomatal ratio (SR, unitless) expressed as ratio of SD_ada_ : SD_aba_, leaf thickness (μm), mesophyll surface area exposed to intercellular air spaces (S_mes_, μm^2^ μm^-2^) and mesophyll cell wall thickness (T_CW_, μm). The details of sample preparation for light and electron microscopy and measurements are presented in Pathare *et al*., (2020a). Light microscopy images of leaf cross sections were used to measure length of mesophyll cell walls exposed to intercellular air spaces (IAS) using 10-15 different fields of view for each leaf (*n* = 3 per species) taken at x 50 and x 100 magnifications. The S_mes_ was calculated from measurements of total length of mesophyll cell walls exposed to IAS and width of section analyzed using equation from Evans *et al*., (1994) with curvature correction factor (F) of 1.34. T_CW_ was measured from TEM micrographs using at least 15 images for each leaf. Values for all the leaf anatomical traits used in current study have been published in Pathare *et al*., 2020a. Values of mean annual precipitation (MAP) for habitats where the C_4_ grasses commonly occur, were obtained as indicated previously in Pathare *et al*., 2020b.

### Enzyme assays and measurement of leaf nitrogen content

Immediately following gas exchange measurements, leaf samples were taken from the same leaf and frozen in liquid nitrogen for enzyme assays. Measurements of carbonic anhydrase (CA), phosphoenolpyruvate carboxylase (PEPC, μmol m^-2^ s^-1^), ribulose-1,5-bisphosphate carboxylase/oxygenase (Rubisco, μmol m^-2^ s^-1^) activities were performed at 25°C as described previously (Sharwood *et al*., 2016; Sonawane & Cousins, 2019; Pathare *et al*., 2020a). CA activities were expressed as first order rate constant (k_CA_, μmol m^-2^ s^-1^ Pa^-1^). In addition to enzyme activities, PEPC’s affinity for HCO^-^_3_ (K_m_, μM HCO_3_^-1^) values were derived using the membrane-inlet mass spectrometer. K_m_ values have been published in Dimario *et al*., 2021. For measuring leaf nitrogen content, leaf samples were taken from the same leaf on which gas exchange measurements were performed. Samples were dried in hot air oven at 60 °C for 72 hours. Leaf nitrogen content was measured using a Eurovector elemental analyzer and expressed on leaf area basis (N_area_, gm^-2^).

### Statistical analyses

Statistical analyses were performed using R software (v 4.1.0, R Foundation for Statistical Computing, Vienna, Austria). Normality and equal variances were tested and when necessary square root or log transformations were used to improve the data homoscedasticity (Zar, 2007). One-way ANOVA with post-hoc Tukey’s test was used to examine differences in leaf-level physiological, anatomical, and biochemical traits among the 16 diverse C_4_ grasses. Results for post-hoc Tukey’s test are given in Table 1 and S1 to S4. For the one-way ANOVA, *P*-values ≤ 0.05 were considered as statistically significant. Results of one-way ANOVA for traits used in the current study are given in Table S1. In addition to the one-way ANOVA, Pearson correlation coefficients were calculated for relationships of percent change in physiological traits (A_net_, g_sw_, g_m_ and TE_i_) with important anatomical and biochemical traits likes T_CW_, S_mes_, SR, SD_ada_, Leaf thickness, PEPC activity, Rubisco activity, k_CA_, K_m_, A_max_ and N_area_. For the trait-to-trait correlations, *P*-values ≤ 0.05 were considered as statistically significant, whereas *P*-values ≤ 0.1 were considered as marginally significant.

To complement the trait-to-trait correlations, we also performed a principal component analysis (PCA), to identify the major axes of variation based on the important anatomical and biochemical traits and the percent change in physiological traits (Fig. S5, Table S2). The R package FactoMineR (Le *et al*., 2008) was used to perform PCA. All traits were scaled during the analysis. The first three principal components (PC) had eigenvalues greater than one (Table S2) and were retained according to Kaiser’s rule (Kaiser, 1960). For each trait, factor loadings greater than 0.5 in absolute value were considered important.

## Results

### CO_2_ response of physiological traits

The 16 C_4_ grasses (listed in Table 1) with previously demonstrated variation in leaf-level anatomical traits and mesophyll conductance (Pathare *et al*., 2020a; Pathare *et al*., 2020b) were chosen to explore potential variation in CO_2_ response of g_m_ and its relation to leaf anatomical and biochemical traits as well as TE_i_. Species differed significantly in CO_2_ response of g_m_ (Fig.1 and Fig. S1) and g_sw_ (Fig. S2). In general, across the 16 C_4_ grasses, g_m_ increased as CO_2_S and C_i_ decreased (Fig.1 and S1) with percent increase at lowest C_i_ ranging from +13% to +250%. Like g_m_, g_sw_ increased with decreasing CO_2_S (Fig. S2). However, unlike g_m_, the magnitude of increase in g_sw_ was lower. Specifically, the percent increase in g_sw_ ranged from about 40-80% when CO_2_S decreased from 34 to 14 Pa. Alternatively, both A_net_ and TE_i_ decreased with decreases in CO_2_S (Fig. S3 and S4). Particularly, A_net_ decreased by about 23-40% and TE_i_ decreased by about 53-64% when CO_2_S decreased from 34 to 14 Pa.

**Figure 1.**
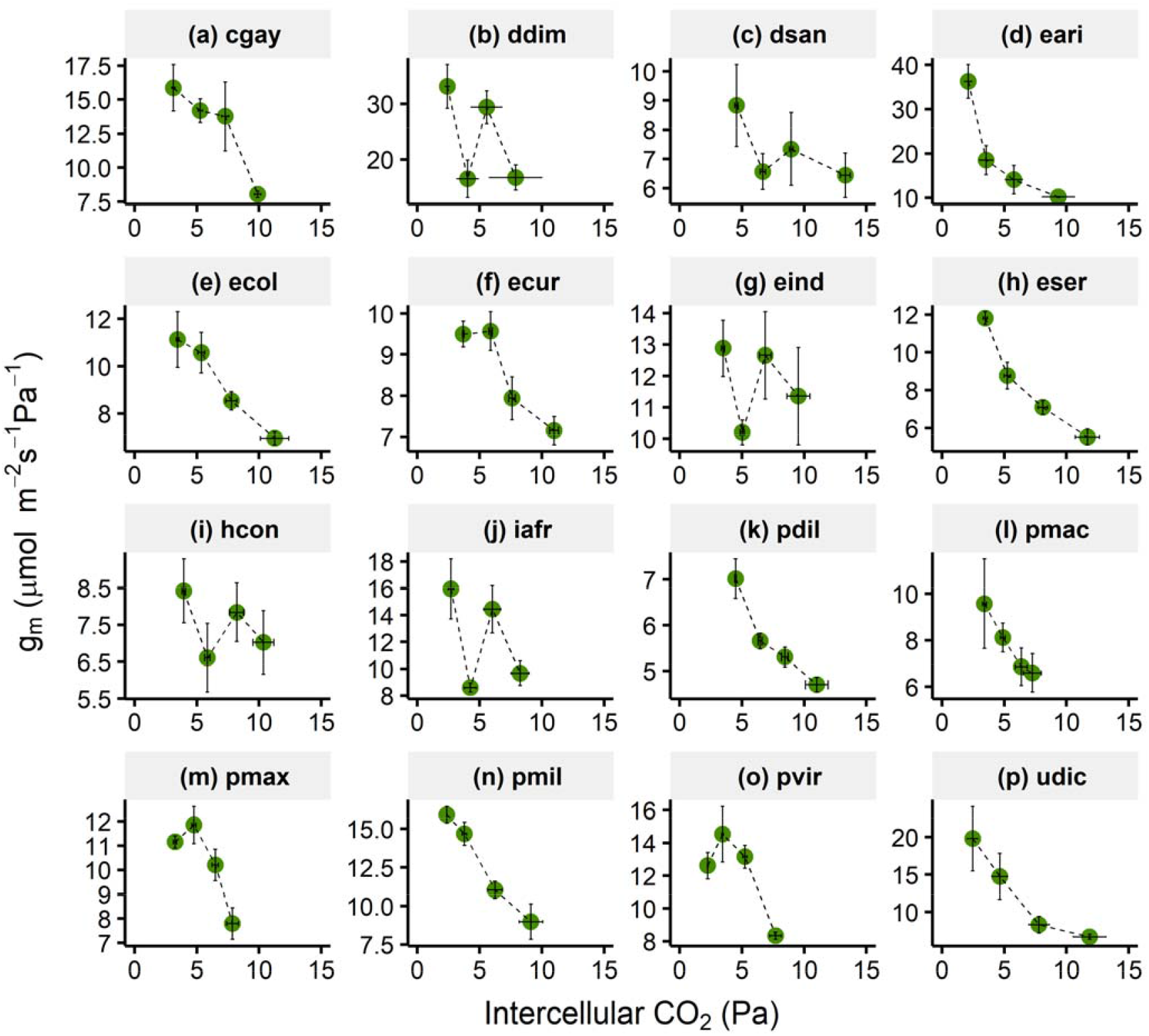
Response of mesophyll conductance (g_m_) to changes in intercellular *p*CO_2_ in 16 diverse C_4_ grasses measured in current study. Measurements were performed at constant light (photosynthetic photon flux density (PPFD) = 1200 μmol m^-2^ s-^1^) and leaf temperature (25°C). Values in each panel represent mean ± SE with *n* = 3-6. Response of g_m_ to CO_2_ S for each species is plotted in separate panel. Species code has been indicated first letter of genera and first three letters of species (see Table 1 for full names of species). *p* values from one-way ANOVA along with Tukey’s letters are shown in Fig. S1.

### Variation in leaf anatomical and biochemical traits

The results of one-way ANOVA suggest that the 16 C_4_ grasses vary significantly (*p* < 0.001, Table 1 and S1, Pathare *et al*., 2020a, DiMario *et al*., 2021) in all the leaf-level anatomical and biochemical traits listed in Table 1. Mesophyll cell wall thickness (T_CW_) showed a significant 2.6-fold variation with values ranging from about 0.08 to 0.21 μm. S_mes_ showed a 2.4-fold variation with values ranging from about 8.5 to 19 μm^2^ μm^-2^. The C_4_ grasses measured here also showed a highly significant variation in stomatal traits like SR and SD_ada_ (*p* < 0.001; Table 1 and S1, Pathare *et al*., 2020a). SR varied by 4.7-fold, with values ranging from 0.5 to 2.4, whereas SD_ada_ showed a 13-fold variation with values ranging from 12 to 160 number mm^-2^. Leaf thickness showed a significant 2-fold variation with values ranging from 120 to 240 μm (*p* < 0.001, Table 1 and S1). Similar to the variation in anatomical traits mentioned above, the biochemical traits (that is leaf-level activities of PEPC, Rubisco and k_CA_) also varied significantly among the 16 C_4_ grasses included in the current study (*p* < 0.001, Table 1 and S1). PEPC activities showed a 7.5-fold variation with values from 96 to 721 μmol m^-2^ s^-1^. Rubisco activities showed a 2.6-fold variation with values from 30 to 80 μmol m^-2^ s^-1^. k_CA_ showed an 18-fold variation with values from 9 to 165 μmol m^-2^ s^-1^ Pa^-1^. PEPC’s affinity for HCO_3_^-^ (*K*_m_) showed a 1.6-fold variation across the 16 C_4_ grasses with values ranging from around 29 to 46 μM HCO_3_^-^ (*p* < 0.001, Table 1 and S1, DiMario *et al*., 2021). There was a 1.7-fold variation in maximum photosynthetic rates (A_max_) with values ranging from ∼25 to 42.5 μmol CO_2_ m^-2^ s^-1^ (*p* < 0.001, Table 1). N_area_ also varied significantly among the grasses (*p* < 0.001, Table 1) with values ranging from 1.24 to 2.26 gm^-2^ (Pathare *et al*., 2020a).

### Relationship of CO_2_ response of g_m_ with CO_2_ response of TE_i_, g_sw_ and A_net_

We investigated the impacts of changes in g_m_ with *p*CO_2_ on corresponding changes in TE_i_, gsw and A_net_ (Fig. 2). For this we calculated percent change in the trait’s value when CO_2_S decreases from 34 to 14 Pa. Since g_m_ and g_sw_ increase with decreases in CO_2_S, percent change for both conductances is reported as percent increase in the manuscript (Figs. 2-5). Alternatively, A_net_ and TE_i_ decrease with decreases in CO_2_S, thus the percent change for these two traits have been reported as percent decrease in the manuscript (Fig. 2 and S6-S8).

**Figure 2.**
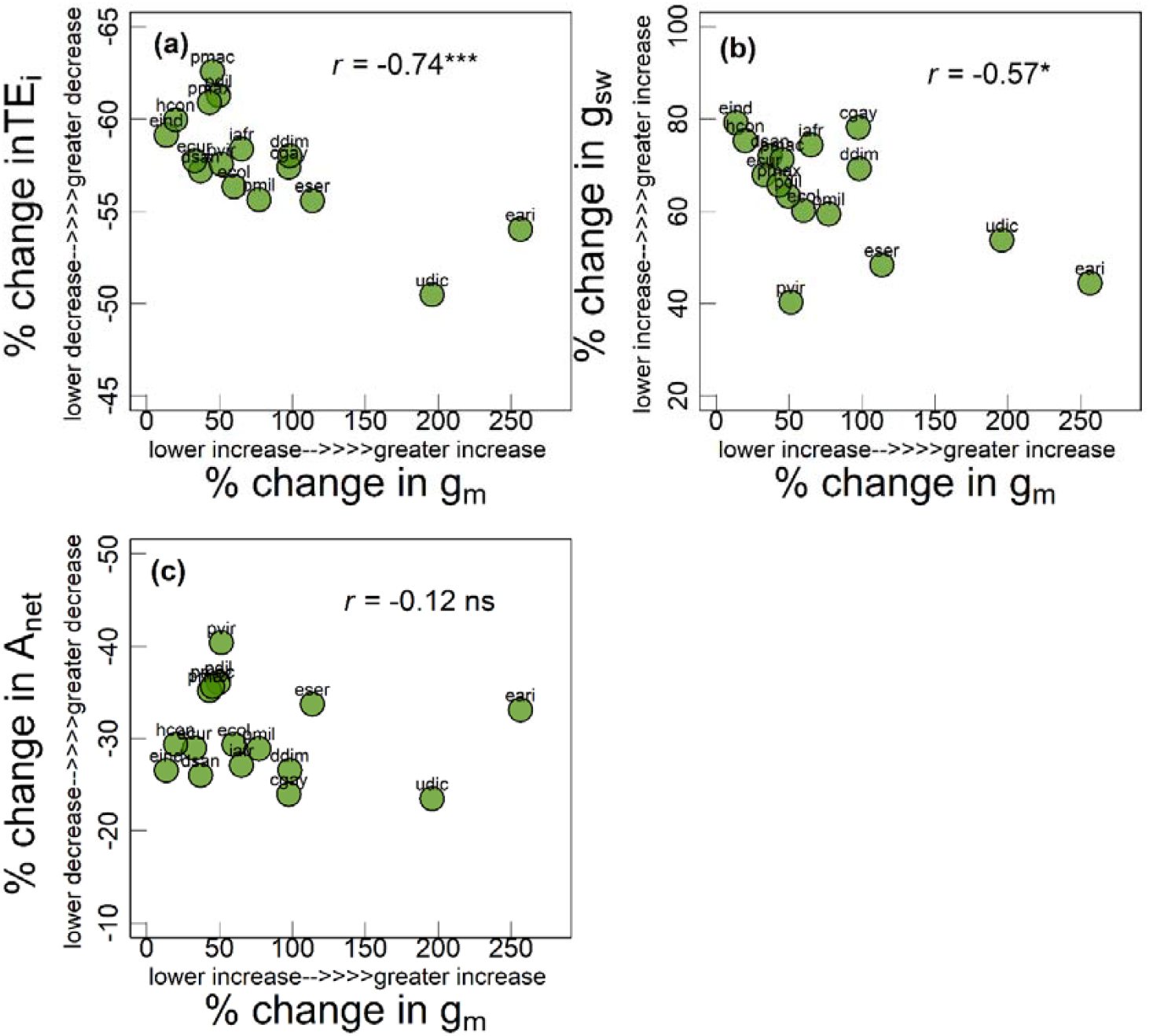
Relationship of percent increase in g_m_ with (a) percent decrease in leaf-level water-use efficiency (TE_i_) expressed as A_net_/g_sw_ (higher negative value indicates greater decrease in TE_i_), (b) percent increase in stomatal conductance to water (g_sw_) and (c) percent decrease in net photosynthetic rates (A_net_) (higher negative value indicates greater decrease in A_net_) for the 16 C_4_ grasses measured in current study. Significance of Pearson correlation coefficients: ^ns^, non-significant, +, marginally significant, *, *p* ≤ 0.05, **, *p* ≤ 0.01 and ***, *p* ≤ 0.001. Each circle represents mean value for each species (*n* = 3-6). Species names are indicated by codes given in Table 1.

Percent change in TE_i_ showed a strong negative correlation with percent change in g_m_ (*r* = -0.74, *p* < 0.01, Fig. 2a). Particularly, C_4_ grasses which were able to achieve a greater increase in g_m_ at lower CO_2_S (for example, udic and eari) showed a lesser decrease in TE_i_ (indicated by a less negative value for TE_i_ in Fig. 2a). Whereas, percent change in g_sw_ showed a strong negative relationship with percent change in g_m_ (*r* = -0.57, *p* = 0.019, Fig. 2b), that is, species which showed a greater increase in g_m_ at lower CO_2_S also showed a lesser increase in g_sw_. For example, species eari showed the greatest increase in g_m_ (250 %) but the lowest increase in g_sw_ (40%). A significant relationship was not observed between percent change in g_m_ and A_net_ (Fig. 2c).

### Relationships of CO_2_ response of physiological traits with anatomical and biochemical traits

Relationship of percent change in g_m_ with anatomical and biochemical traits are shown in Figs. 3 and 4. Percent change in g_m_ was negatively related to T_CW_ (*r* = -0.60, *p* = 0.01, Fig. 3a), that is, species with thinner cell walls showed greater increases in g_m_ at lower CO_2_S. Whereas, percent change in g_m_ showed a statistically significant positive relationship with SR (*r* = 0.54, *p* = 0.03, Fig. 3c) and leaf thickness (*r* = 0.52, *p* < 0.05, Fig. 3e) and a non-significant but positive relationship with S_mes_ (*r* = 0.25, *p* > 0.1, Fig. 3b) and SD_ada_ (*r* = 0.34, *p* > 0.1, Fig. 3d). Percent change in g_m_ also showed a non-significant but positive relationship with biochemical traits like PEPC activity (*r* = 0.29, *p* > 0.1, Fig. 4a) and *K*_m_ (*r* = 0.26, *p* > 0.1, Fig. 4d) and a significant positive relationship with Rubisco activity (*r* = 0.46, *p* = 0.07, Fig. 4b) and *k*_CA_ (*r* = 0.42, *p* = 0.1, Fig. 4c).

**Figure 3.**
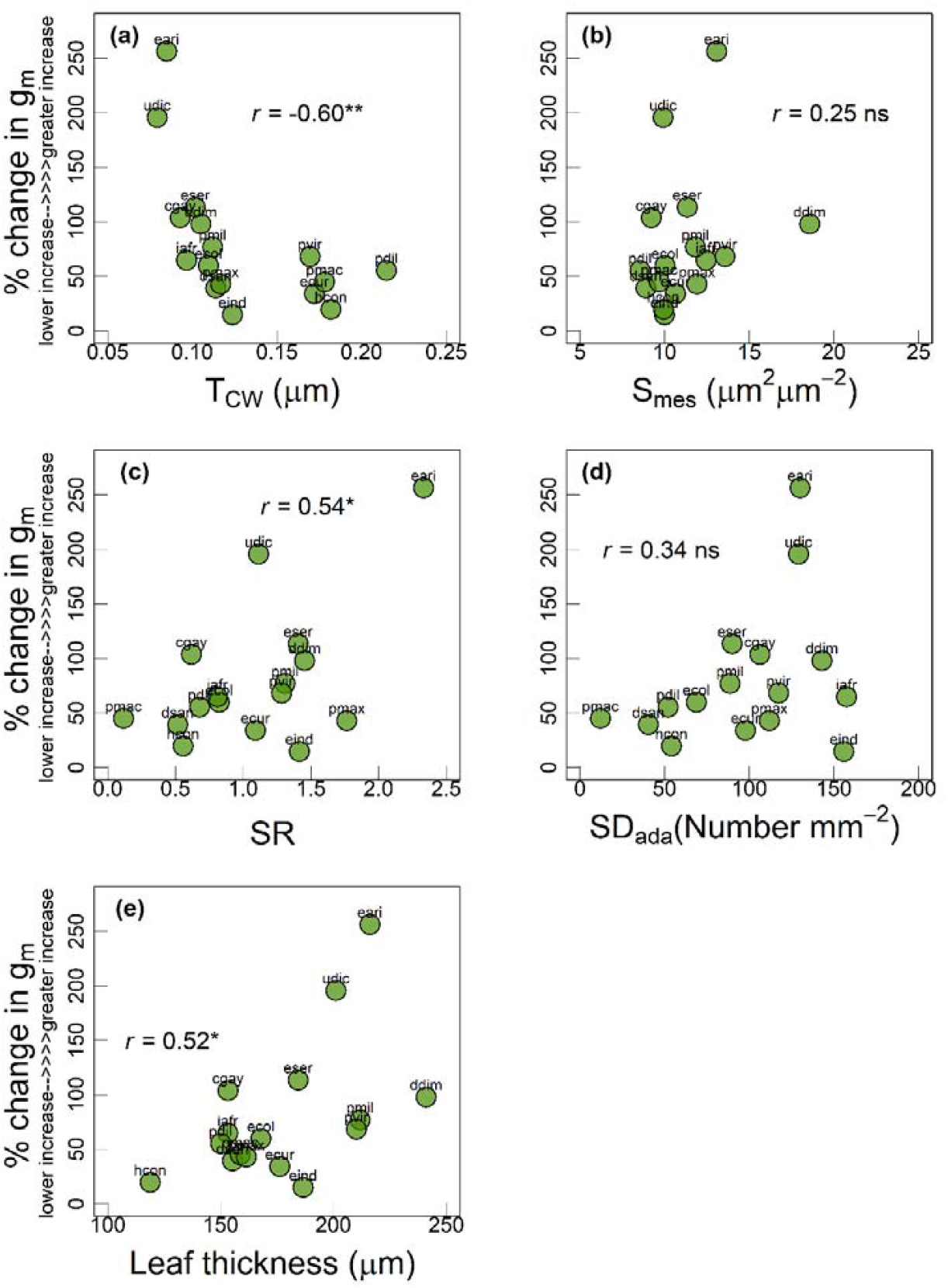
Relationship of percent increase in g_m_ with (a) mesophyll cell wall thickness (T_CW_) (b) mesophyll surface area exposed to intercellular air spaces (S_mes_) (c) stomatal ratio (SR), (d) stomatal density adaxial (SD_ada_) and (e) leaf thickness among the 16 C_4_ grasses measured in current study. Significance of Pearson correlation coefficients: ^ns^, non-significant, +, marginally significant, *, *p* ≤ 0.05, **, *p* ≤ 0.01 and ***, *p* ≤ 0.001. Each circle represents mean value for each species (*n* = 3-6). Species names are indicated by codes given in Table 1.

**Figure 4.**
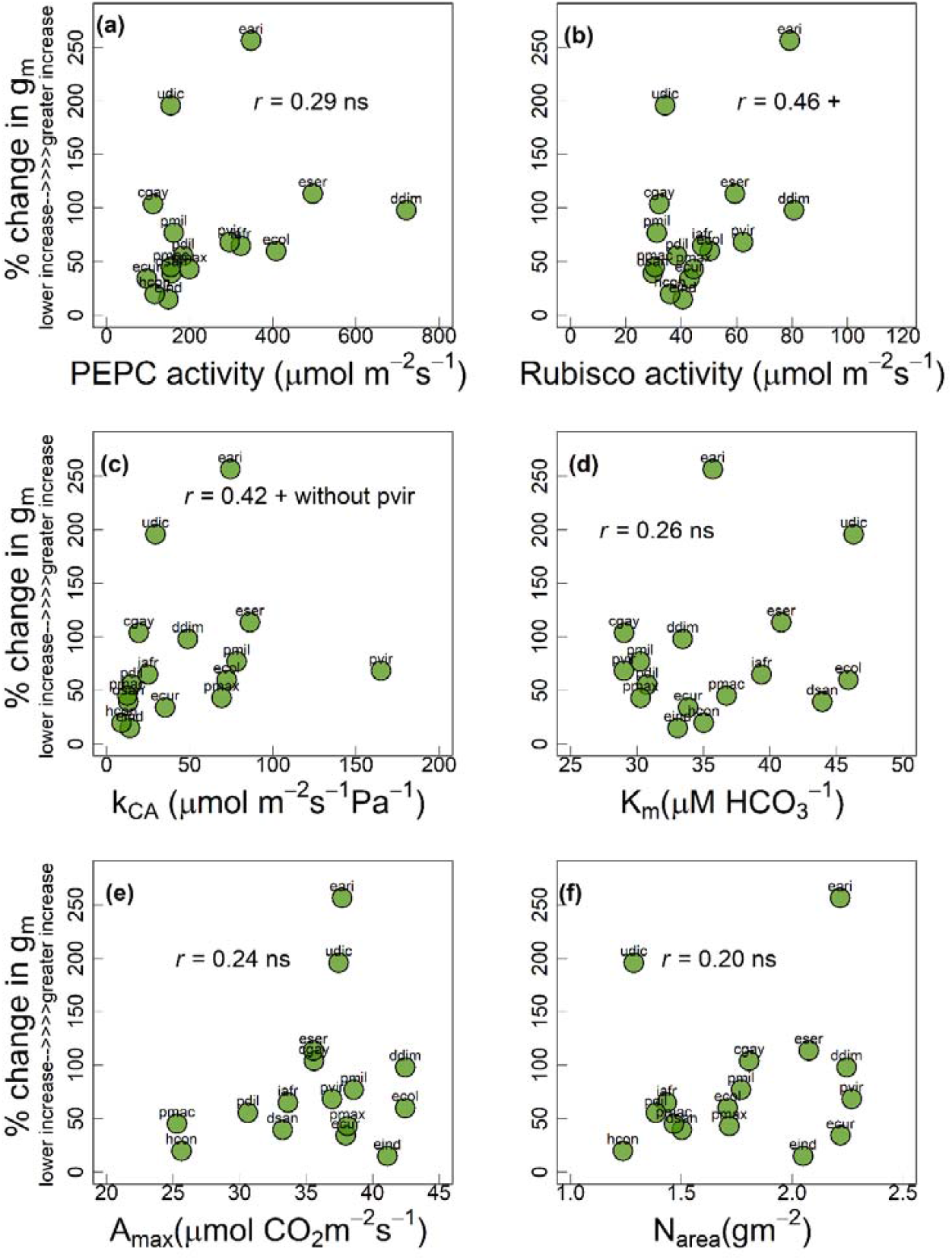
Relationship of percent increase in g_m_ with (a) PEPC activity (b) Rubisco activity (c) CA activity expressed as k_CA_, (d) PEPC’s affinity for HCO_3_^-^ (K_m_), (e) maximum photosynthetic capacity (A_max_) and (f) leaf N content (N_area_) among the 16 C_4_ grasses measured in current study. Significance of Pearson correlation coefficients: ^ns^, non-significant, +, marginally significant, *, *p* ≤ 0.05, **, *p* ≤ 0.01 and ***, *p* ≤ 0.001. Each circle represents mean value for each species (*n* = 3-6). Species names are indicated by codes given in Table 1.

To account for effects of anatomical traits, particularly T_CW_ which was strongly related to CO_2_ response of C_4_ g_m_, and biochemical traits on CO_2_ response of g_m_, we derived the ratio of T_CW_ with PEPC activity, Rubisco activity, k_CA_, *K*_m_, A_max_ and N_area_ and investigated the relationship of percent change in g_m_ with these ratios (Fig. 5). Percent change in g_m_ showed significant negative relationship with T_CW_/PEPC (*r* = -0.52, *p* = 0.05, Fig. 5a), T_CW_/Rubisco (*r* = -0.59, *p* < 0.05, Fig. 5b), T_CW_/*K*_m_ (*r* = -0.55, *p* < 0.05, Fig. 5c) and T_CW_/*k*_CA_ (*r* = -0.47, *p* = 0.06, Fig. 5d). In summary, C_4_ grasses with thinner cell walls and greater enzyme activities or *K*_m_ were able to achieve greater increase in g_m_ at lower CO_2_S. Also, in contrast to percent change in g_m_, percent change in g_sw_ was positively related to T_CW_ and negatively related to enzyme activities (as indicated by PCA Fig. S5 and Table S1). Thus, grasses with lower T_CW_ but greater A_max_, N_area_, enzyme activities and greater values for K_m_ showed a lesser increase in g_sw_ at lower CO_2_S.

**Figure 5.**
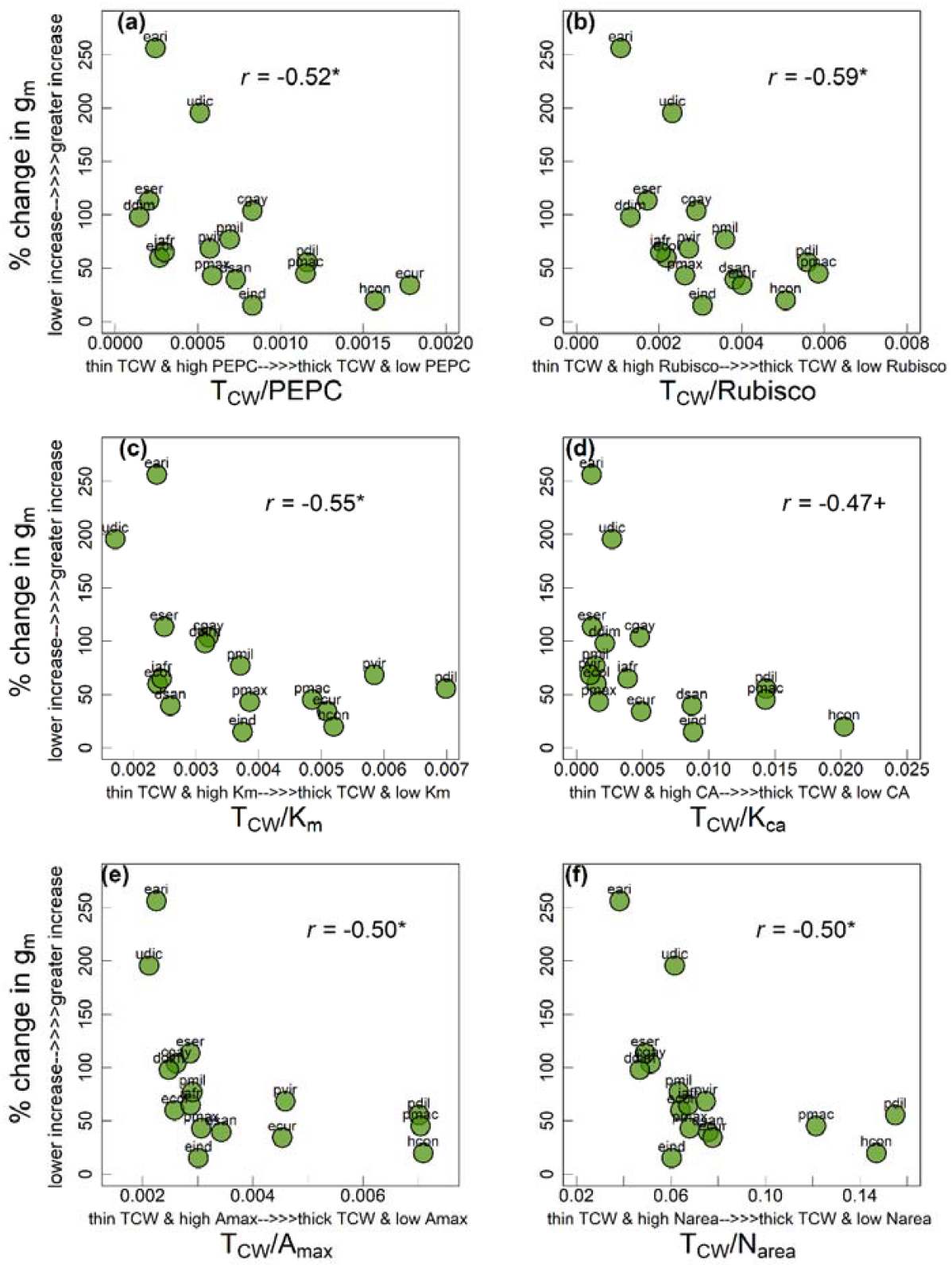
Relationship of percent increase in g_m_ with ratio of mesophyll cell wall thickness (T_CW_) to (a) PEPC activity (b) Rubisco activity (c) PEPC’s affinity for HCO_3_^-^ (K_m_), (d) CA activity expressed as k_CA_, (e) maximum photosynthetic rates (A_max_) and (f) leaf N content (N_area_) among the 16 C_4_ grasses measured in current study. Significance of Pearson correlation coefficients: ^ns^, non-significant, +, marginally significant, *, *p* ≤ 0.05, **, *p* ≤ 0.01 and ***, *p* ≤ 0.001. Each circle represents mean value for each species (*n* = 3-6). Species names are indicated by codes given in Table 1.

Like percent change in g_m_, we also investigated the relationship of percent change in TE_i_ with above mentioned anatomical (Fig. S6), biochemical traits (Fig. S7) and ratios of T_CW_ with biochemical traits (Fig. S8). Percent change in TE_i_ showed a statistically significant positive relationship with T_CW_ (*r* = 0.68, *p* < 0.01, Fig. S6a). Whereas percent change in TE_i_ showed a statistically significant negative relationship with SR (*r* = -0.41, *p* = 0.1, Fig. S6c) and a non-significant negative relationship with SD_ada_ (*r* = -0.40, *p* > 0.1, Fig. S6d). In terms of its relationship with biochemical traits, percent change in TE_i_ showed a non-significant negative relationship with PEPC activity (*r* = -0.16, *p* > 0.1, Fig. S7a), Rubisco activity (*r* = -0.21, *p >* 0.1, Fig. S7b), *k*_CA_ (*r* = -0.27, *p* > 0.1, Fig. S7c) and significant negative relationship with *K*_m_ (*r* = -0.48, *p* = 0.05, Fig. S7d). Furthermore, percent change in TE_i_ showed a significant positive relationship with T_CW_/PEPC (*r* = 0.46, *p* = 0.07, Fig. S8a), T_CW_/Rubisco (*r* = 0.62, *p* = 0.01, Fig. S8b), T_CW_/*K*_m_ (*r* = 0.67, *p* < 0.01, Fig. S8c) and T_CW_/*k*_CA_ (*r* = 0.60, *p* = 0.013, Fig. S8d). In summary, C_4_ grasses with thinner cell walls but greater SR, leaf thickness, S_mes_, SD_ada_, enzyme activities and *K*_m_ showed a lesser decrease in TE_i_ at lower CO_2_S.

### Principal Component Analysis

To complement the trait-to-trait correlations, we also performed a PCA using the important leaf-level anatomical and biochemical traits and percent changes in g_m_, A_net_, g_sw_ and TE_i_ and mean annual precipitation. The PCA was performed on leaf traits from 16 C_4_ grasses, where the first four axes with eigenvalues ≥ 1 were retained for analysis (Table S2). The first two major axes (PC1 and PC2) of leaf trait variation along with the average position of C_4_ grasses in PC1-PC2 space are presented in Figure S5. PC1 explains about 44 % of total variation in the C_4_ grasses and has a positive association with traits like SD_ada_, S_mes_, SR, leaf thickness, PEPC activity, Rubisco activity, k_CA_, A_max_, N_area_ and percent change in g_m_ and TE_i_. Whereas PC1 showed negative association with T_CW_ and percent change in g_sw_ and mean annual precipitation. Thus, PC1 delineates the C_4_ grasses into those with greater CO_2_ response of g_m_ (higher PC1; greater enzyme activity and lower T_CW_) and those with lower CO_2_ response of g_m_ (lower PC1; lower enzyme activity and greater T_CW_). Alternatively, PC2 explains about 18 % of the total variation in the C_4_ grasses and is positively associated with *K*_m_ and percent change in A_net_ and TE_i_ and negatively associated with T_CW_ and k_CA_. PC2 tends to separate the C_4_ grasses into those showing greater *K*_m_ values, lower T_CW_ and lesser decrease in A_net_ and TE_i_ at low CO_2_S.

## Discussion

Changes in g_m_ in response to CO_2_ have been well studied for C_3_ species. In general C_3_-g_m_ increases under short-term decreases in *p*CO_2_ (Flexas *et al*., 2007; Bunce, 2010; Douthe *et al*., 2011), though not always (von Caemmerer & Evans, 1991; Loreto *et al*., 1992; Tazoe *et al*., 2009). The main candidates suggested to affect the CO_2_ response of C_3_-g_m_ include chloroplast movement and hence changes in Sc, changes in activities of carbonic anhydrase (Evans *et al*., 2009; Momayyezi & Guy, 2017) and the facilitation effect of CO_2_-permeable aquaporins (Uehlein *et al*., 2008; Flexas *et al*., 2012; Kaldenhoff, 2012). However, the C_3_-g_m_ response to *p*CO_2_ may also result from systematic errors associated with use of methods like gas exchange, chlorophyll fluorescence and discrimination against ^13^C or the use of oversimplified models (Pons *et al*., 2009; Yin & Struik, 2009; Tholen & Zhu, 2011; Gu & Sun, 2014). For instance, the apparent response of C_3_-g_m_ to short-term changes in *p*CO_2_ has partly been attributed to changes in photorespiratory and non-photorespiratory release of CO_2_ to the total CO_2_ pool in the leaf particularly under low *p*CO_2_. Ignoring these CO_2_ pools while estimating C_3_-g_m_ also overlooks the effects of spatial distribution of mitochondria and chloroplast on pathlength of CO_2_ movement. Inaccurate measurements of day respiration or estimates of the Rubisco fractionation factor in the Δ^13^C method and measurements conducted at low [O_2_] have also been suggested as potential sources of artefacts in determining the C_3_-g_m_ response to *p*CO_2_ (Pons *et al*., 2009; Gu & Sun, 2014).

Alternatively, in C_4_ species, the photorespiratory and non-photorespiratory release of CO_2_ is relatively low and may not contribute significantly to estimates of g_m_ even at low *p*CO_2_ (Cousins et al., 2020). Also, in current study we estimated g_m_ in diverse C_4_ grasses by using a method based on modelling of Δ^18^O that considers the contribution of respiratory fluxes(Ogee *et al*., 2018). Comparing many diverse C_4_ species, as done in current study, using a method that is not subject to same limitations as those previously used for C_3_ species, is a reasonable approach to identify CO_2_ responses of C_4_-g_m_. Here, we discuss the CO_2_ response of g_m_ for 16 diverse C_4_ grasses and its relationships with leaf anatomical and biochemical traits.

### CO_2_ response of g_m_ varied among diverse C_4_ grasses

Very few studies have investigated the CO_2_ response of g_m_ in C_4_ species, primarily maize and *Setaria viridis* (Osborn *et al*., 2017; Kolbe & Cousins, 2018; Ubierna *et al*., 2018). In general, these studies reported an increase in C_4_-g_m_ with a decrease in *p*CO_2_. Similarly, we demonstrate that g_m_ increases with a decrease in *p*CO_2_ across the 16 diverse C_4_ grasses. However, compared to previous studies, the magnitude of increase in g_m_ varied greatly across the 16 C_4_ grasses (+13% to +250%; Fig. 1). In the following sections we discuss the potential factors that could explain this variability in CO_2_ response of C_4_-g_m_.

### CO_2_ response of C_4_-g_m_ is related with T_CW_ and photosynthetic capacity

g_m_ in C_3_ and C_4_ species is constrained by several anatomical and biochemical parameters like leaf thickness, stomatal density, T_CW_, S_mes_, S_c_, CA activities and aquaporins (Cousins *et al*., 2020; Evans, 2021). However, the potential implication of leaf anatomy and biochemistry for CO_2_ response of g_m_ has not been studied for C_4_ species. Most of the anatomical parameters remain unchanged under short-term changes in environmental conditions like *p*CO_2_ (Evans *et al*., 2009; Terashima *et al*., 2011). Hence, the only anatomical trait suggested to influence CO_2_ response of g_m_ is chloroplast movement which can affect g_m_ by changing S_c_ (Terashima *et al*., 2006; Tholen *et al*., 2008; Carriqui *et al*., 2018). However, chloroplast movement is unlikely to explain the variability of CO_2_ response of C_4_-g_m_ observed in the current study, because S_mes_ and not S_c_ is a more accurate determinant of C_4_-g_m_ (Barbour *et al*., 2016; Pathare *et al*., 2020a). In contrast to Sc, S_mes_ should remain unchanged under short-term changes in *p*CO_2_. Hence, though we observed a positive relationship between CO_2_ response of C_4_-g_m_ and S_mes_ (Fig. 3b), S_mes_ may provide only a partial explanation for this variable response. Other anatomical traits like stomatal density and leaf thickness also showed a positive relationship with CO_2_ response of C_4_-g_m._ However, like S_mes_, stomatal density and leaf thickness will also remain unchanged under short-term changes in *p*CO_2_ and may provide only a partial explanation for variable CO_2_ response of C_4_-g_m_.

T_CW_ could account for >50% of the total resistance to CO_2_ diffusion inside leaves (Evans *et al*., 2009). Hence, we investigated the influence of T_CW_ on variability of CO_2_ response of C_4_-g_m_. In general, C_3_ species with relatively lower T_CW_ show greater g_m_ (Onoda *et al*., 2017; Veromann-Jürgenson *et al*., 2017; Evans, 2021). We did not observe a strong relationship between g_m_ and T_CW_ for the C_4_ grasses under ambient CO_2_ levels (34 Pa) (Pathare *et al*., 2020a). However, the CO_2_ response of C_4_-g_m_ was related to T_CW_ in current study (Fig. 3a). Particularly, C_4_ grasses with relatively lower T_CW_ showed greater increase in g_m_ at low *p*CO_2_. It’s unclear why g_m_ of C_4_ grasses with lower T_CW_ is more responsive to changes in *p*CO_2_. Like other anatomical traits, T_CW_ is unlikely to change under short-term changes in *p*CO_2_ and cannot be the reason for observed variable CO_2_ response of C_4_-g_m_. However, T_CW_ may still provide partial explanation for variability in CO_2_ response of C_4_-g_m_. Resistances to CO_2_ diffusion through liquid phase are greater compared to gaseous phase (Evans *et al*., 2009). CO_2_ must dissolve in the water-filled pores of the mesophyll cell walls and then diffuse to the plasma membrane and eventually to the site of CO_2_ fixation. Because cell walls represent a significant proportion of liquid phase resistance (Evans *et al*., 2009), C_4_ species with lower T_CW_ may have the potential to achieve a greater change in g_m_ in response to changing *p*CO_2_. However, the influence of lower T_CW_ on variability of CO_2_ response of C_4_-g_m_ may also be augmented by the facilitation effect of CO_2_ transporting aquaporins and proteins of photosynthetic machinery that are involved in drawdown and fixation of CO_2_ (Parkhurst, 1994; Wright *et al*., 2004; Evans *et al*., 2009).

The role of CO_2_-permeable aquaporins in enhancing g_m_ has been well characterized in C_3_ species (Uehlein *et al*., 2008). Only recently, it has been demonstrated that overexpressing a CO_2_-permeable aquaporin in plasma membrane of *Setaria viridis* (C_4_ grass), can enhance C_4_-g_m_ (Ermakova *et al*., 2021). We did not investigate the role of aquaporins in affecting CO_2_ response of C_4_-g_m_. However, we observed a greater CO_2_ response of g_m_ in C_4_ grasses with relatively lower T_CW_. This indicates that g_m_ in plants with lower T_CW_ may be more influenced by aquaporins (Evans, 2021). There is still a need to further investigate role of aquaporins in variable CO_2_ response of C_4_-g_m_.

Here, we investigated the relationship between CO_2_ response of C_4_-g_m_ and photosynthetic capacity as indicated by A_max_, N_area_ and activities of key C_4_ photosynthetic enzymes (PEPC, CA and Rubisco). CO_2_ response of C_4_-g_m_ showed a positive, but non-significant, relationship with A_max_, N_area_ and PEPC, Rubisco and CA activities (Fig. 4). Thus, photosynthetic capacity alone is not strongly related to CO_2_ response of C_4_-g_m_. Instead, greater photosynthetic capacity in combination with relatively lower T_CW_ are associated with a greater CO_2_ response of C_4_-g_m_ (Fig. 5). Lower T_CW_ may have decreased the resistance to movement of CO_2_ into the mesophyll cells (Evans, 2021; Flexas *et al*., 2021). Whereas greater photosynthetic capacity may have increased the demand for CO_2_ and hence the necessity of maintaining a greater CO_2_ supply through an increase in g_m_ at low *p*CO_2_ (Wright *et al*., 2004; Evans *et al*., 2009). Furthermore, the enzyme CA catalyzes the conversion of CO_2_/HCO_3_^-^ in the cytosol and ensures sufficient HCO_3_^-^ substrate supply to PEPC (Studer *et al*., 2014; DiMario *et al*., 2018). Because CO_2_ diffuses much faster in liquid phase in HCO_3_^-^ form, greater CA activities could have maintained a rapid supply of HCO_3_^-^ substrate to PEPC and further enhanced g_m_ at low *p*CO_2_ in C_4_ grasses with lower T_CW_.

We also investigated the relationship between the CO_2_ response of C_4_-g_m_ and K_m_ - a kinetic constant indicating PEPC’s affinity for HCO_3_^-^ (DiMario *et al*., 2021). In general, lower K_m_ values (or high affinity of PEPC for HCO_3_^-^) are expected to provide a selective advantage by maintaining high rates of C_4_ photosynthesis, particularly under conditions like drought when CO_2_ availability is low due to restricted stomatal conductance. Here, we observed that K_m_ alone did not show a strong relationship with CO_2_ response of C_4_-g_m_ (Fig. 4d). However, after accounting for T_CW_, we observed that T_CW_/K_m_ shows a significant negative relationship with CO_2_ response of C_4_-g_m_ (Fig. 5c), wherein, species with relatively lower T_CW_ and higher K_m_ (or lower affinity of PEPC for HCO_3_^-^) exhibited a greater CO_2_ response of g_m_ (Fig. 5c). This contrasts the general expectation and could be explained by the lower T_CW_ leading to higher *g*_m_ and with the corresponding greater CA activities lead to higher HCO_3_^-^ in mesophyll cells under low C_i_ conditions. The higher HCO_3_^-^ concentration in the mesophyll cells of species with lower T_CW_ and greater CA can reduce the selective pressure on PEPC for lower K_m_ values.

### Relationship of CO_2_ response of g_m_ with CO_2_ response of g_sw_ and TE_i_

Previously, we demonstrated that C_4_-g_m_ is positively related to TE_i_ under ambient *p*CO_2_ (Pathare *et al*., 2020a). Our current study further suggests that for C_4_ grasses, g_m_ may also influence TE_i_ under short-term changes in *p*CO_2_. In general, both g_m_ and g_sw_ increased (Fig. 1 and S2) and TE_i_ decreased at low *p*CO_2_ (Fig. S4). However, the magnitude of increase in g_m_ at low *p*CO_2_ was greater (values from 13% to 250%) compared to increase in g_sw_ (values from 40% to 80%). Also, C_4_ grasses showing greatest increase in g_m_ at low *p*CO_2_ also showed the lowest increase in g_sw_ (Fig. 2b). Consequently, though TE_i_ decreased at low *p*CO_2_, the decrease was less in the species showing greater CO_2_ response of g_m_ (Fig.2a). Such species with greater CO_2_ response of g_m_ could have a particular advantage, in terms of maintaining TE_i_ under low CO_2_ conditions like drought, compared to species whose g_m_ was less responsive to changes in *p*CO_2_. Indeed, we observed that C_4_ grasses with a greater CO_2_ response of g_m_ are generally adapted to relatively drier habitats (Fig. S5).

## Conclusions

We demonstrated that C_4_-g_m_ increases with decrease in *p*CO_2_ and the magnitude of this increase in g_m_ varies greatly among the 16 diverse C_4_ grasses. Also, CO_2_ response of C_4_-g_m_ is a composite trait that seems to be influenced by many leaf anatomical and biochemical parameters. The greatest increase in g_m_ at low *p*CO_2_ was observed in C_4_ grasses with lower T_CW_ and greater photosynthetic capacities. These C_4_ grasses with a greater CO_2_ response of g_m_ were also able to maintain their TE_i_ under low *p*CO_2_, which may be advantageous under low CO_2_ conditions like drought. Our study advances understanding about CO_2_ response of g_m_ in diverse C_4_ species and identifies the key leaf anatomical and biochemical traits related to this response. This understanding is essential for improving C_4_ photosynthetic models (von Caemmerer, 2021) and in attempts to improve water-use efficiency of C_4_ crops through modification of g_m_ (von Caemmerer & Furbank, 2016).

## Acknowledgements

This work was supported by the Division of Chemical Sciences, Geosciences, and Biosciences, Office of Basic Energy Sciences, Department of Energy (grant no. DE-SC0001685), Office of Biological and Environmental Research in the DOE Office of Science (DE-SC0018277), and the National Science Foundation (Major Research Instrumentation grant no. 0923562). We are grateful to Dr Nerea Ubierna, Joseph Crawford and Dr Balasaheb Sonawane for their valuable inputs on methods of estimation of mesophyll conductance in C4 species. We are also grateful to the Core Facility Center “Cell and Molecular Technologies in Plant Science” of Komarov Botanical Institute (St.-Petersburg, Russia) and Franceschi Microscopy and Imaging Center at Washington State University (Pullman, USA) for the use of its facilities and staff assistance. We would also like to thank Charles A. Cody for help in plant growth management. The authors declare that they have no conflicts of interest.

## Author contributions

V.S.P and A.B.C designed the experiment. V.S.P, R.J.D and N.K performed the measurements and analyzed the data. V.S.P interpreted the data and led the writing with constructive inputs from R.J.D, N.K and A.B.C.

