## Supplemental for "Mesophyll conductance response to short-term changes in CO_2_ is related to leaf anatomy and biochemistry in diverse C_4_ grasses"

Orchid ID: 0000-0001-6220-7531 (V.S.P.); 0000-0002-5056-6868 (R.J.D); 0000-0002-7507-4560 (N.K.); 0000-0003-2424-714X (A.B.C)

**Summary**

- Mesophyll CO_2_ conductance (g_m_) in C_3_ species responds to short-term (minutes) changes in environment potentially due to changes in some leaf anatomical and biochemical properties and due to measurement artifacts. Compared to C_3_ species, there is less information about g_m_ responses to short-term changes in environment conditions like *p*CO_2_ across diverse C_4_ species and the potential determinants of these responses.
- Using 16 diverse C_4_-grasses we investigated the response of g_m_ to short-term changes in CO_2_ and how this response related to the leaf anatomical and biochemical traits.
- For all the measured C_4_-grasses g_m_ increased as CO_2_ decreased; however, the percent change in g_m_ varied (+13% to +250%) and significantly related to percent changes in leaf transpiration efficiency (TE_i_). The percent increase in g_m_ was highest in grasses with thinner mesophyll cell walls and greater leaf nitrogen, activities of phosphoenolpyruvate carboxylase (PEPC), Rubisco and carbonic anhydrase, and a higher affinity of PEPC for bicarbonate substrate.
- Our study demonstrates that CO_2_ response of g_m_ varies greatly across diverse C_4_-grasses and identifies the key leaf anatomical and biochemical traits related to this variation. These findings have implications for improving C_4_ photosynthetic models, and in attempts to improve TE_i_ through manipulation of g_m_.

**Table S1** Results of one-way ANOVA with species as main effects for all the leaf-level anatomical and biochemical traits measured for 16 C_4_ grasses in current study.

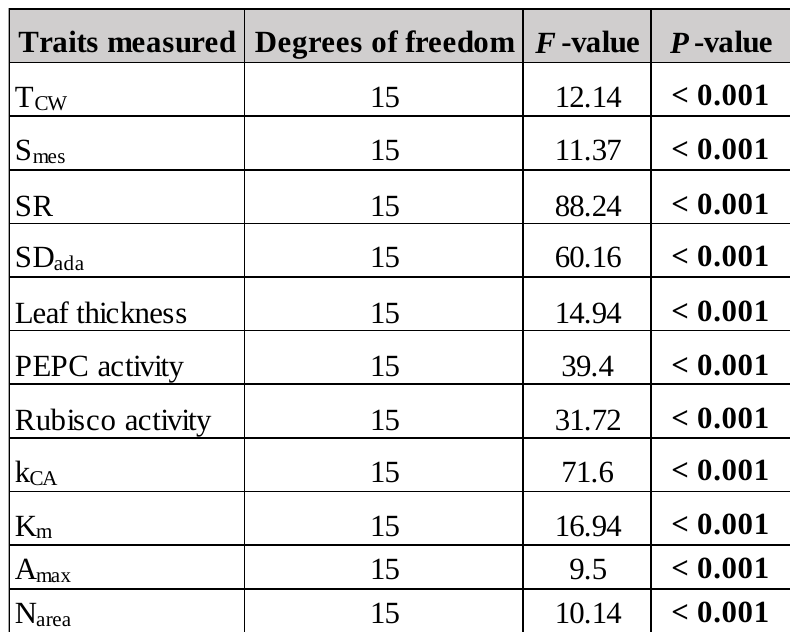

Statistically significant *P*-values (≤ 0.05) are highlighted in bold. Numerator degrees of freedom and *F*-values (*F* test statistic) are also shown. Mesophyll cell wall thickness (T_CW_), total mesophyll cell surface area exposed to intercellular air space per unit of leaf surface area (S_mes_), stomatal ratio (SR), adaxial stomatal density (SD_ada_), Carbonic anhydrase activity expressed as first-order rate constant (k_CA_), phosphoenolpyruvate carboxylase activity (PEPC), PEPC’s affinity for HCO^-^_3_ (K_m_), maximum photosynthetic capacity (A_max_) and leaf N content (N_area_).

**Tables S2** Component loadings for important leaf-level traits determined on 16 C_4_ grasses.

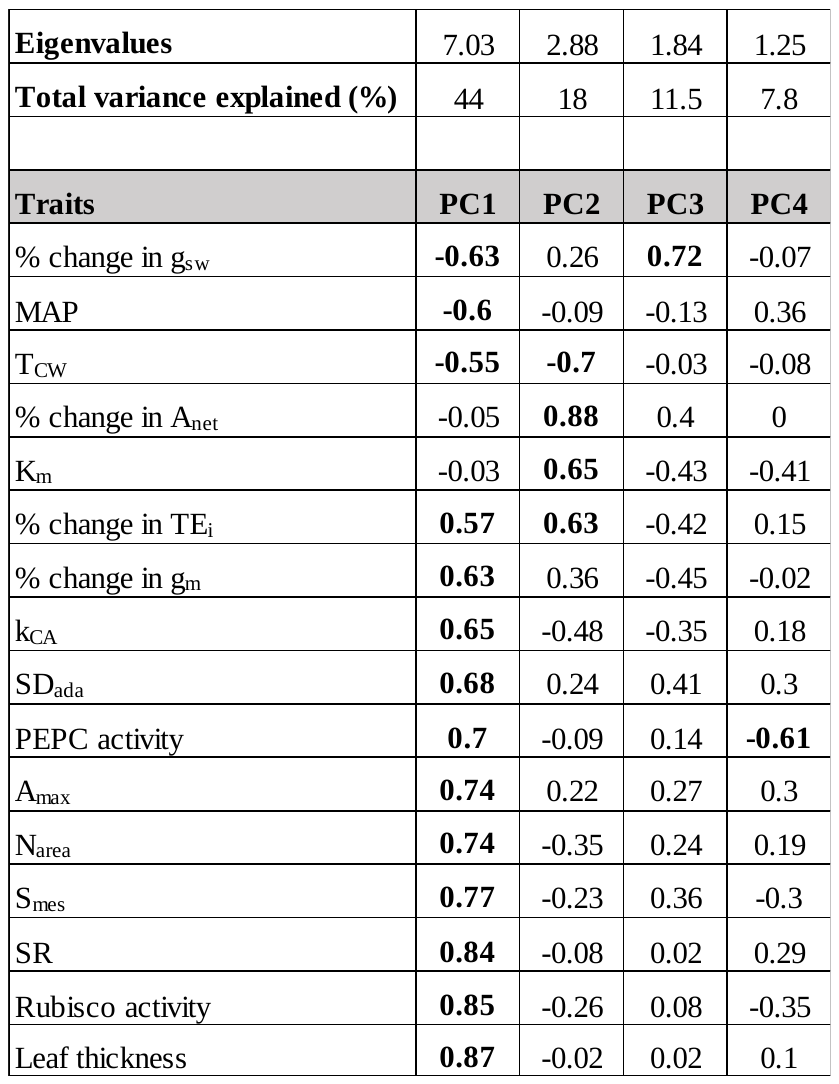

Eigenvalues and total variance explained (%) by first four principal components (PC1, PC2, PC3, PC4) is shown. The first four PCs have eigenvalues ≥ 1 and hence are retained. Component loadings greater than 0.5 in absolute value are shown in bold face. Mean annual precipitation (MAP), Mesophyll cell wall thickness (T_CW_), intrinsic water use efficiency (TE_i_), stomatal conductance to water vapor diffusion (g_sw_), net photosynthetic rate per unit of leaf surface area (A_net_), total mesophyll cell surface area exposed to intercellular air space per unit of leaf surface area (S_mes_), mesophyll conductance to CO_2_ diffusion estimated by Ogee *et al*., 2018 method (g_m_), stomatal ratio (SR), adaxial stomatal density (SD_ada_), Carbonic anhydrase activity expressed as first-order rate constant (k_CA_), phosphoenolpyruvate carboxylase activity (PEPC), PEPC’s affinity for HCO^-^_3_ (K_m_), maximum photosynthetic capacity (A_max_) and leaf N content on area basis (N_area_).

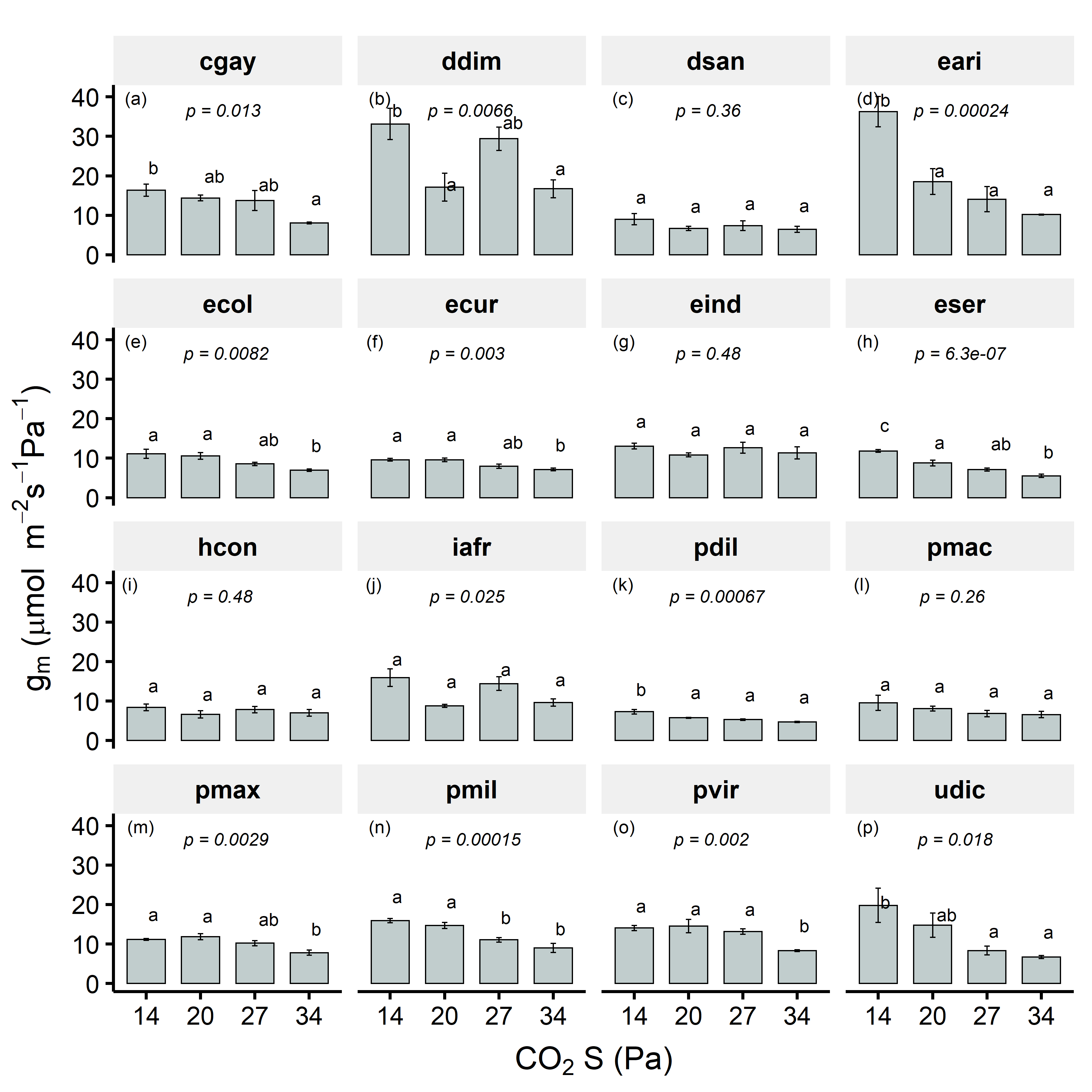

**Figure S1. Response of mesophyll conductance to CO_2_ (g_m_) to changes in CO_2_ sample (CO_2_ S) in 16 diverse C_4_ grasses measured in current study.** Measurements were performed at constant light (photosynthetic photon flux density (PPFD) = 1200 µmol m^-2^ s-^1^) and leaf temperature (25°C). Values in each panel represent mean ± SE with *n* = 3-6. Response of g_sw_ to CO_2_ S for each species is plotted in separate panel. Species code has been indicated first letter of genera and first three letters of species (see Table 1 for full names of species). *p* values from one-way ANOVA along with Tukey’s letters are shown.

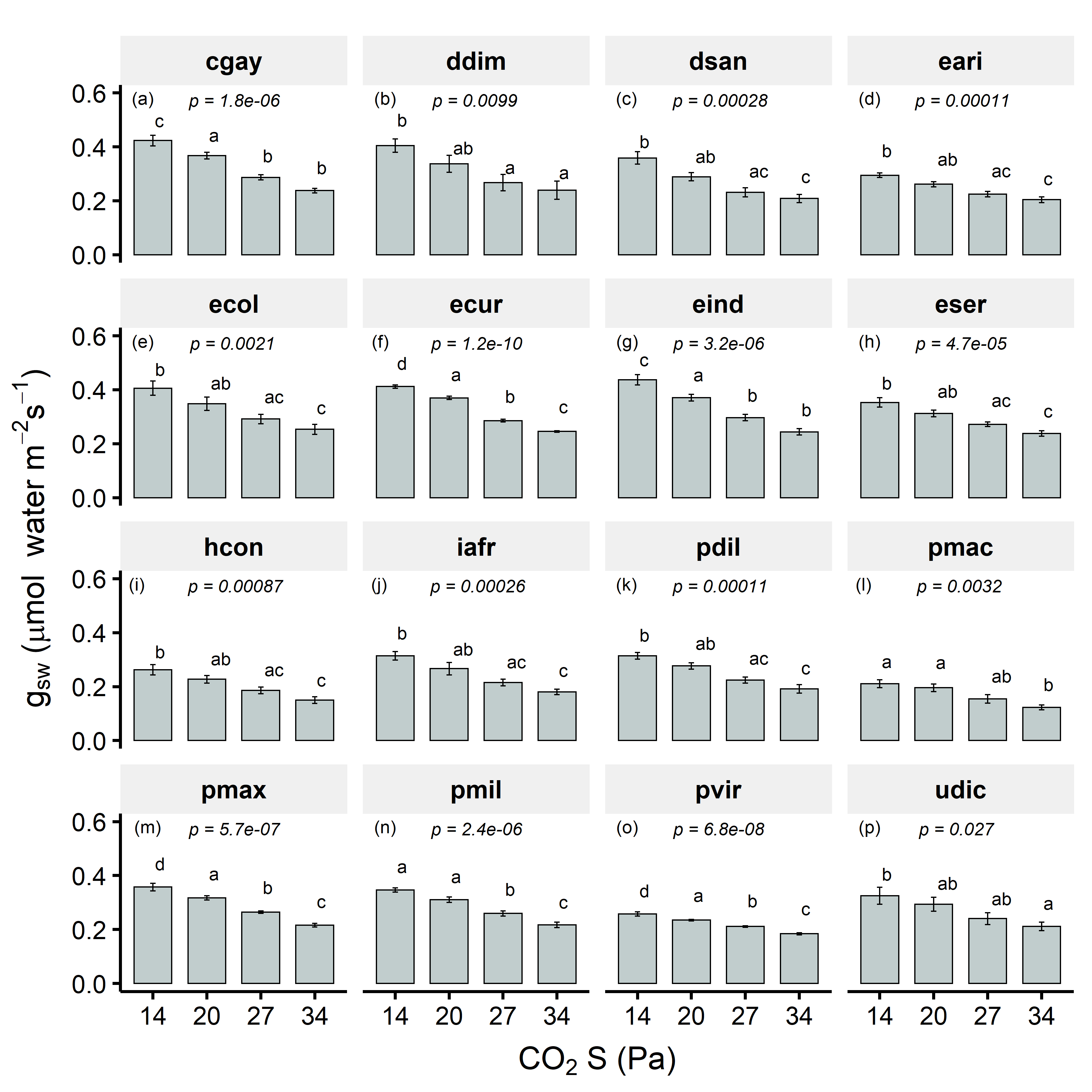

**Figure S2. Response of stomatal conductance to water (g_sw_) to changes in CO_2_ sample (CO_2_ S) in 16 diverse C_4_ grasses measured in current study.** Measurements were performed at constant light (photosynthetic photon flux density (PPFD) = 1200 µmol m^-2^ s-^1^) and leaf temperature (25°C). Values in each panel represent mean ± SE with *n* = 3-6. Response of g_sw_ to CO_2_ S for each species is plotted in separate panel. Species code has been indicated first letter of genera and first three letters of species (see Table 1 for full names of species). *p* values from one-way ANOVA along with Tukey’s letters are shown.

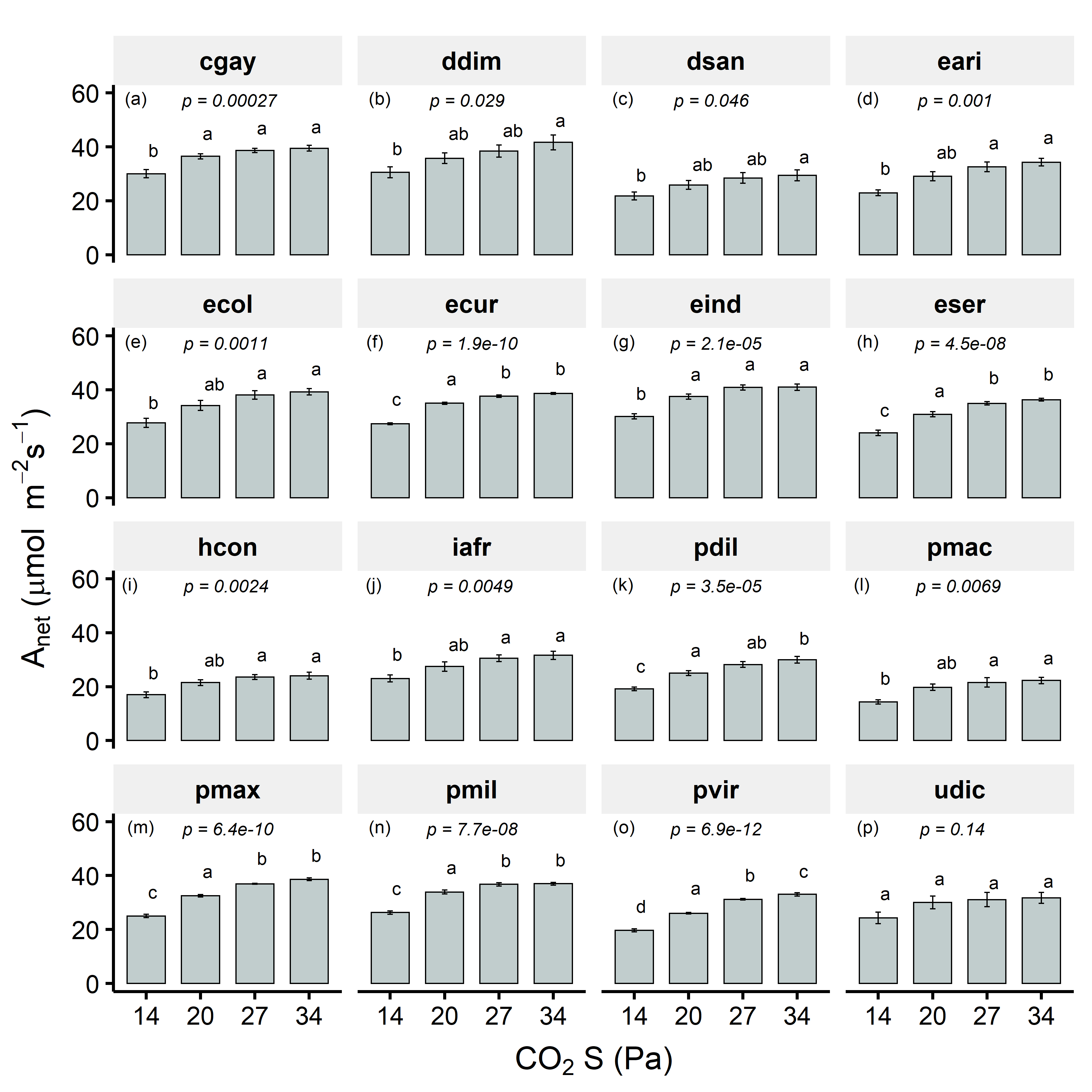

**Figure S3. Response of net CO_2_ assimilation rates (A_net_) to changes in CO_2_ sample (CO_2_ S) in 16 diverse C_4_ grasses measured in current study.** Measurements were performed at constant light (photosynthetic photon flux density (PPFD) = 1200 µmol m^-2^ s-^1^) and leaf temperature (25°C). Values in each panel represent mean ± SE with *n* = 3-6. Response of A_net_ to CO_2_ S for each species is plotted in separate panel. Species code has been indicated first letter of genera and first three letters of species (see Table 1 for full names of species). *p* values from one-way ANOVA along with Tukey’s letters are shown.

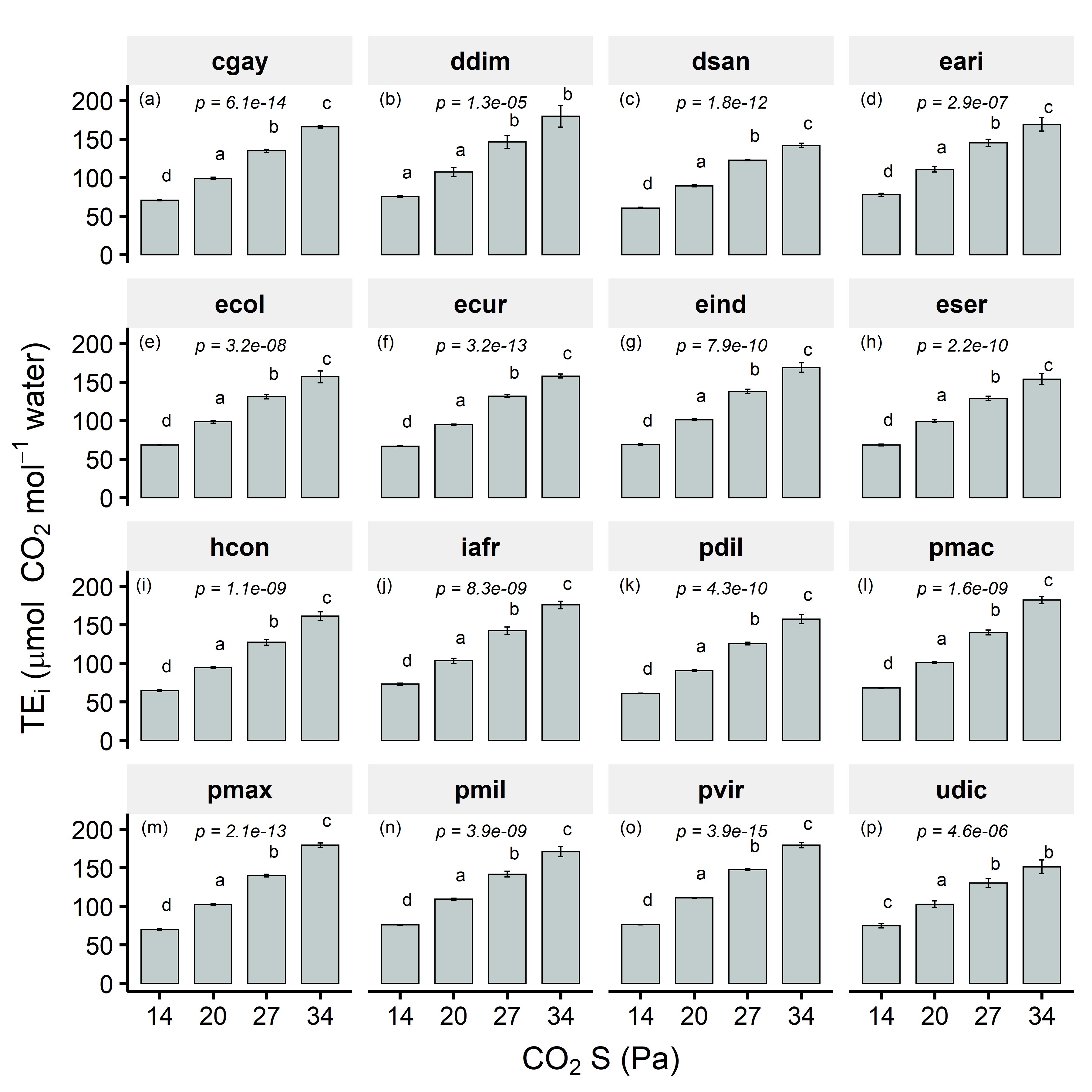

**Figure S4. Response of leaf-level water-use efficiency (TE_i_ = A_net_/g_sw_) to changes in CO_2_ sample (CO_2_ S) in 16 diverse C_4_ grasses measured in current study.** Measurements were performed at constant light (photosynthetic photon flux density (PPFD) = 1200 µmol m^-2^ s-^1^) and leaf temperature (25°C). Values in each panel represent mean ± SE with *n* = 3-6. Response of WUE to CO_2_ S for each species is plotted in separate panel. Species code has been indicated first letter of genera and first three letters of species (see Table 1 for full names of species). *p* values from one-way ANOVA along with Tukey’s letters are shown.

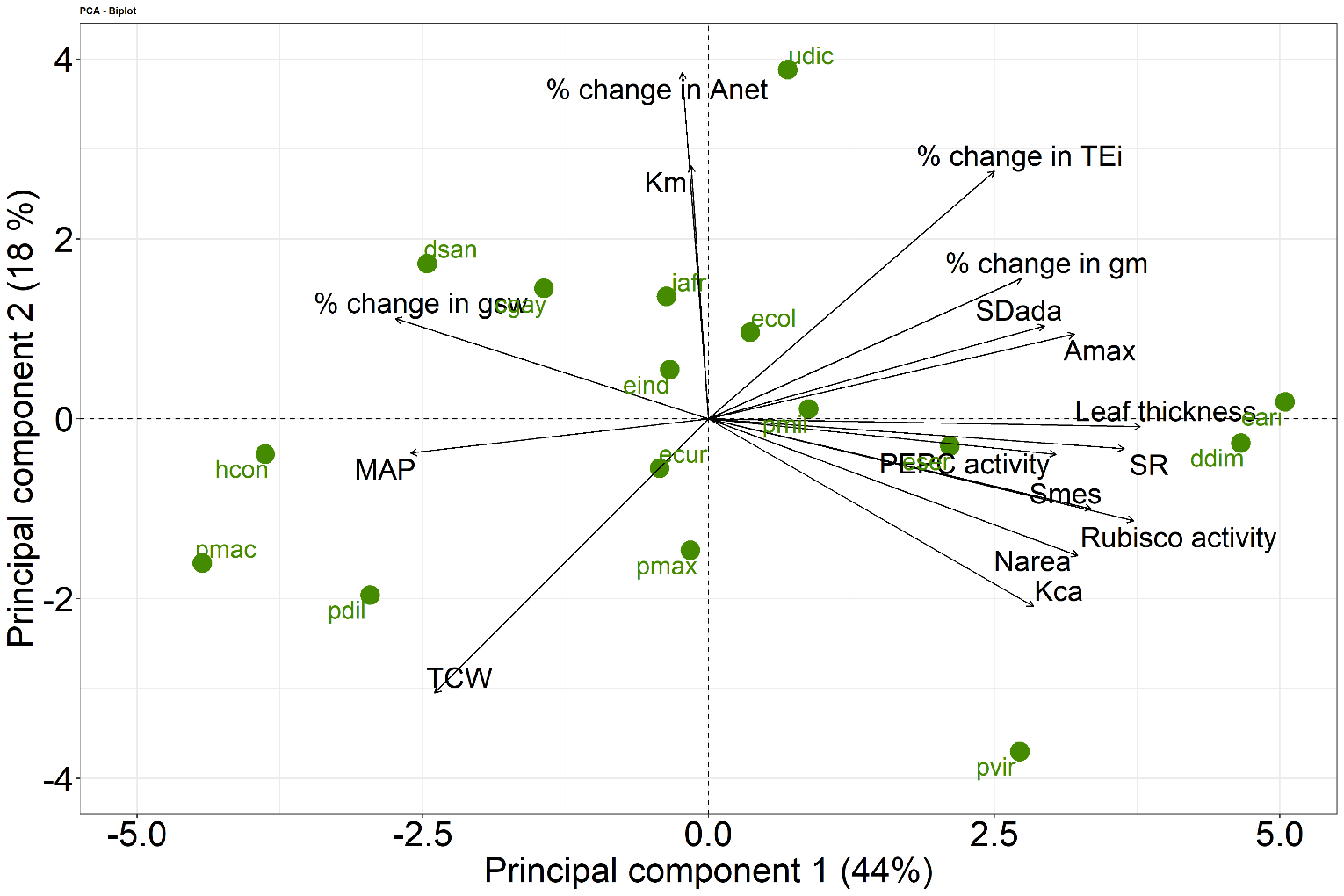

**Figure S5.** PCA biplot showing major axes of variation in important leaf-level anatomical and biochemical traits and percent change (increase or decrease) in response to CO_2_ in physiological traits like g_m_, A_net_, g_sw_ and TEi for the 16 diverse C_4_ grasses measured in current study. Eigenvalues and factor loadings for first three principal components (PC) are shown in Table S1. The arrows are the vectors showing the correlation (across the C_4_ grasses) between a trait and the PCs. Position of species in principal component space is shown in blue circles. Species names correspond to the description in Table 1. Values for MAP and anatomical traits were obtained from Pathare *et al*., 2020a, b, whereas values for K_m_ were obtained from DiMario *et al*., 2021. Points are mean values with *n* = 3-6 per species (mean ± SE values are given in Table 1). g_m_, mesophyll conductance to CO_2_ diffusion; SD_ada_, adaxial stomatal density; S_mes_, total mesophyll cell surface area exposed to intercellular air space per unit of leaf surface area; SR, stomatal ratio; N_area_, leaf N content expressed on area basis; A_max_, maximum photosynthetic rates at saturating light and *p*CO_2_; PEPC, phosphoenolpyruvate carboxylase; Rubisco, ribulose-1,5-bisphosphate carboxylase/oxygenase; Kca, activity of Carbonic anhydrase expressed as first order rate constant; K_m_, PEPC’s affinity for HCO_3_^-^; A_net_, net CO_2_ assimilation rates; g_sw_, stomatal conductance to water; TE_i_, leaf-level water-use efficiency (A_net_/g_sw_); T_CW_, mesophyll cell wall thickness; MAP, mean annual precipitation.

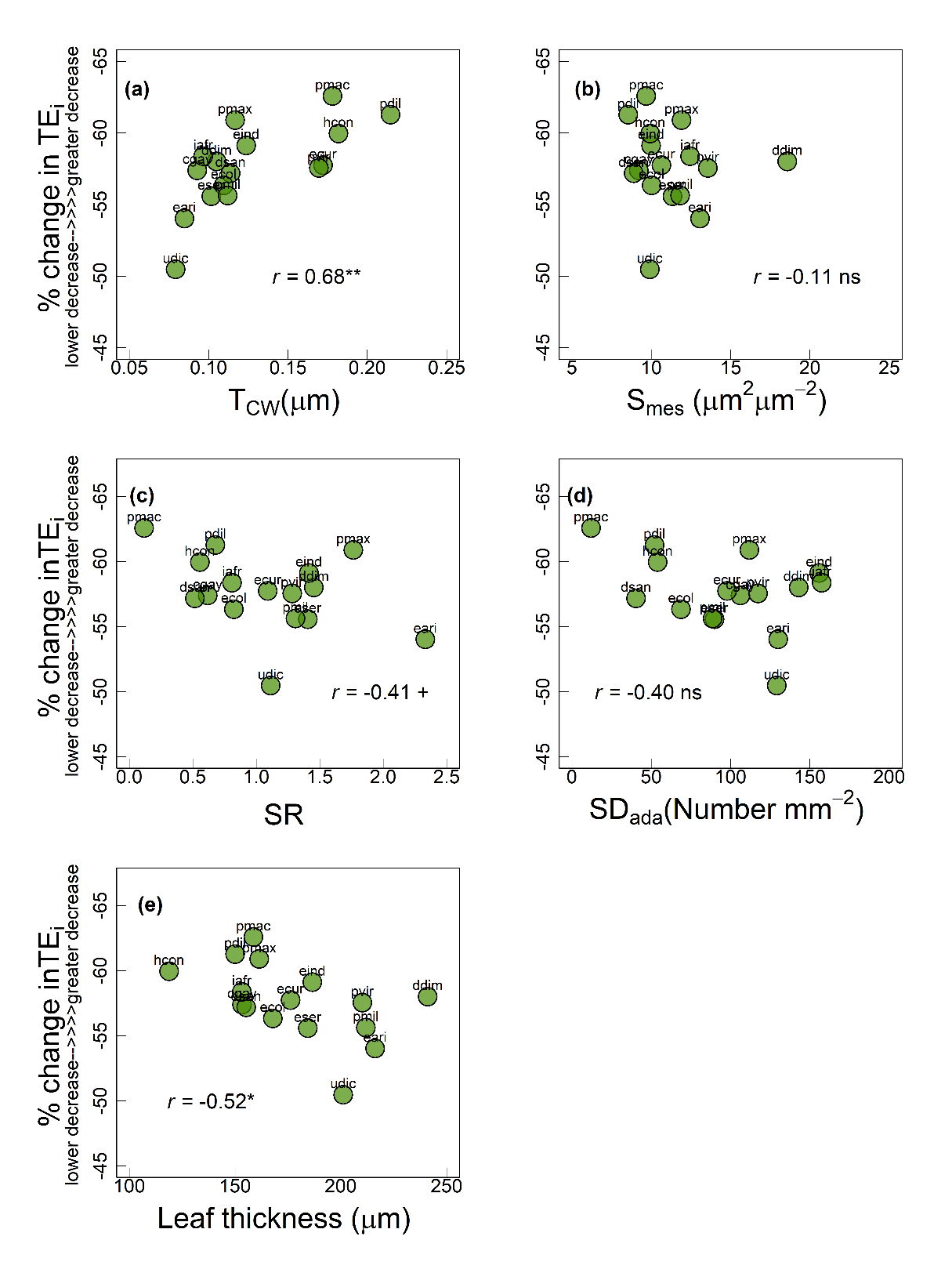

**Figure S6.** Relationship of percent decrease in TE_i_ (higher negative value indicates greater decrease in TE_i_) with (a) mesophyll cell wall thickness (T_CW_) (b) mesophyll surface area exposed to intercellular air spaces (S_mes_) (c) stomatal ratio (SR) and (d) stomatal density adaxial (SD_ada_) among the 16 C_4_ grasses measured in current study. Significance of Pearson correlation coefficients: ^ns^, non-significant, +, marginally significant, *, *p* ≤ 0.05, **, *p* ≤ 0.01 and ***, *p* ≤ 0.001. Each circle represents mean value for each species (*n* = 3-5). Species names are indicated by codes given in Table 1.

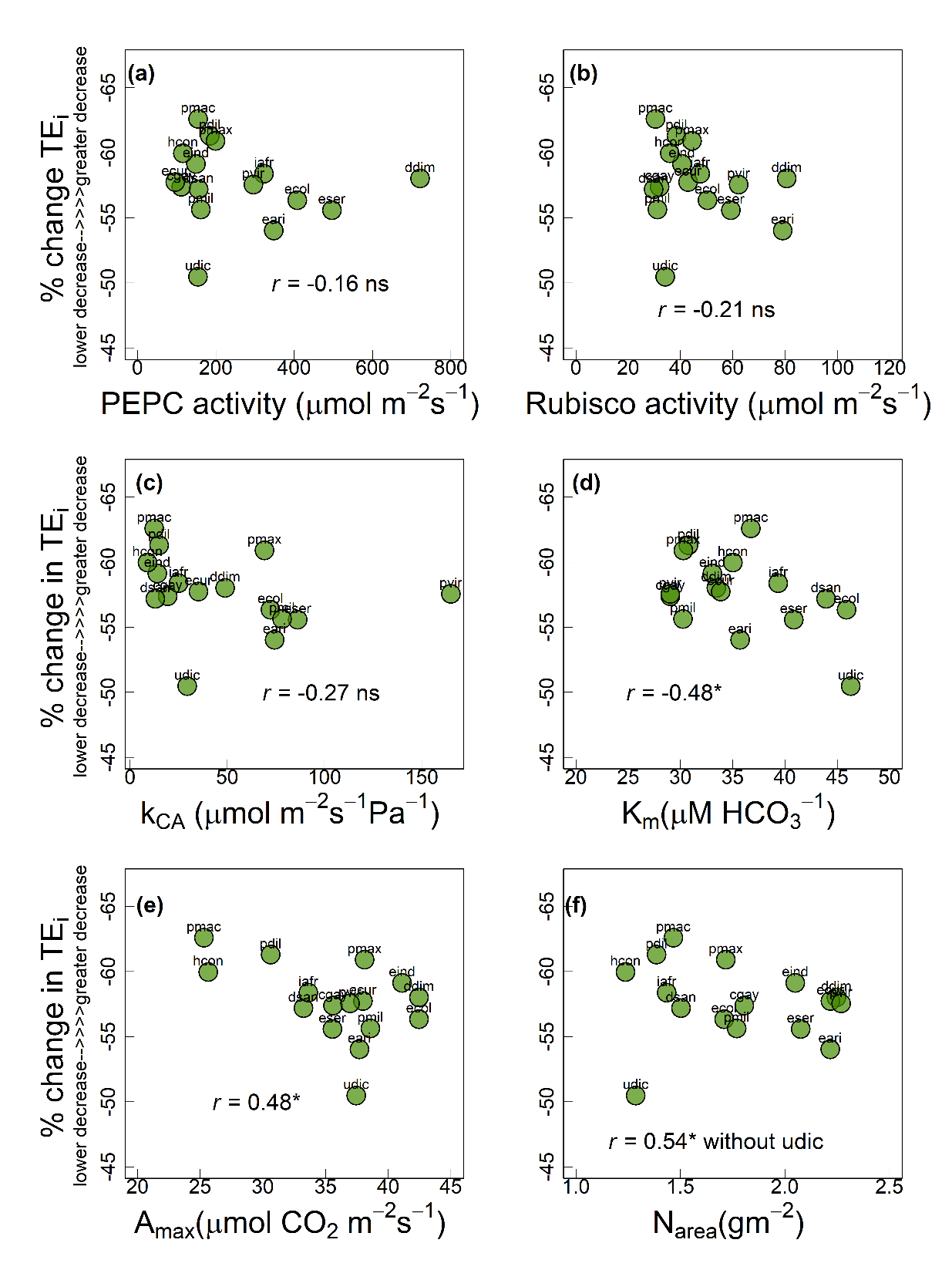

**Figure S7.** Relationship of percent decrease in TE_i_ (higher negative value indicates greater decrease in TE_i_) with (a) PEPC activity (b) Rubisco activity (c) CA activity expressed as k_CA_, (d) PEPC’s affinity for HCO_3_^-^ (K_m_), (e) maximum photosynthetic capacity (A_max_) and (f) leaf N content (N_area_) among the 16 C_4_ grasses measured in current study. Significance of Pearson correlation coefficients: ^ns^, non-significant, +, marginally significant, *, *p* ≤ 0.05, **, *p* ≤ 0.01 and ***, *p* ≤ 0.001. Each circle represents mean value for each species (*n* = 3-6). Species names are indicated by codes given in Table 1.

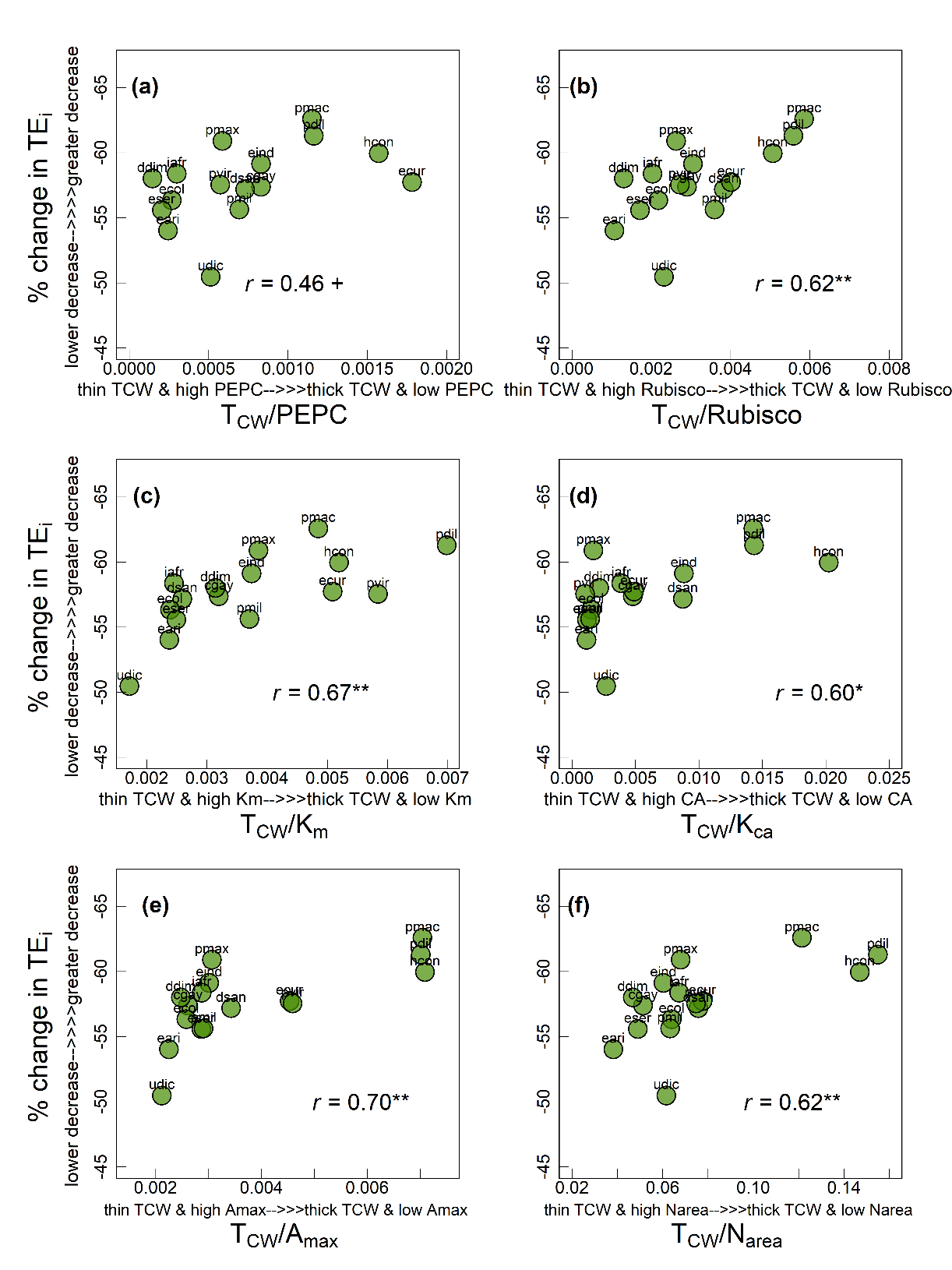

**Figure S8.** Relationship of percent decrease in TE_i_ with ratio of mesophyll cell wall thickness (T_CW_) to (a) PEPC activity (b) Rubisco activity (c) PEPC’s affinity for HCO_3_^-^ (K_m_), (d) CA activity expressed as k_CA_ (e) maximum photosynthetic capacity (A_max_) and (f) leaf N content (N_area_) among the 16 C_4_ grasses measured in current study. Significance of Pearson correlation coefficients: ^ns^, non-significant, +, marginally significant, *, *p* ≤ 0.05, **, *p* ≤ 0.01 and ***, *p* ≤ 0.001. Each circle represents mean value for each species (*n* = 3-5). Species names are indicated by codes given in Table 1.
